# Flexible recruitment of memory-based choice representations by human medial-frontal cortex

**DOI:** 10.1101/809673

**Authors:** Juri Minxha, Ralph Adolphs, Stefano Fusi, Adam N. Mamelak, Ueli Rutishauser

## Abstract

Decisions in complex environments rely on flexibly utilizing past experience as required by context and instructions^1^. This process depends on the medial frontal cortex (MFC) and the medial temporal lobe (MTL)^2-5^, but it remains unknown how these structures jointly implement flexible memory retrieval^6,7^. We recorded single neurons in MFC and MTL while human subjects switched^8^ between making memory- and categorization-based decisions^9,10^. Here we show that MFC rapidly implements changing task demands by utilizing different subspaces of neural activity during different types of decisions. In contrast, no effect of task demands was seen in the MTL. Choices requiring memory retrieval selectively engaged phase-locking of MFC neurons to field potentials in the theta-frequency band in the MTL. Choice-selective neurons in MFC signaled abstract yes-no decisions independent of behavioral response modality (button press or saccade). These findings reveal a novel mechanism for flexibly and selectively engaging memory retrieval^11-14^ and show that unlike perceptual decision-making^15^, memory-related information is only represented in frontal cortex when choices require it.

## Introduction

Behavior in complex environments requires decisions that flexibly combine stimulus representations with context, goals, and memory. Two key aspects of such cognitive flexibility are the ability to selectively utilize relevant information depending on task demands, and to retrieve information from memory when needed. The neural mechanisms that underlie flexible decisions are beginning to be understood in the case of perceptual decision-making ^10,15,16^, with evidence for both early gating, mediated by top-down attention ^17^, and late selection of relevant features in prefrontal cortex ^15^. In contrast, little is known about the mechanisms of decisions that depend in addition also on associated category knowledge and memory, an ability that humans exhibit ubiquitously. In particular, it remains unknown how memory retrieval is selectively engaged when decision-relevant information needs to be actively searched for in memory ^13,18,19^.

The medial frontal cortex (MFC) is critical for complex behavior, and registers cognitive conflict, errors, and choice outcomes ^20-22^. It supports flexible decision-making in two ways: i) by representing task sets ^5,23^ and context ^24^, and ii) by selectively engaging memory retrieval through functional interactions with the amygdala and hippocampus ^3,7,25-28^ (henceforth referred to jointly as the medial temporal lobe, MTL). A mechanism that facilitates such MFC-MTL interactions is phase-locking of MFC activity to oscillations in the MTL, a mode of interaction that has been extensively investigated in rodents during spatial behavior ^6,29,30^ and fear conditioning ^31,32^, but whose broader function remains poorly understood ^33^, particularly in humans. Similarly, findings from human neuroimaging indicate that the MFC is involved in memory search ^2,3,11,12,19,34^, but it remains unknown what features of decisions and context are represented in human MFC, whether MTL-dependent memory retrieval selectively engages synchrony between the two structures, and whether synchrony can be engaged dynamically when required. Our lack of knowledge about the nature of representations in human MFC stands in stark contrast to the patent behavioral ability of humans to flexibly recruit memory processes in everyday life ^8,14^ and to our detailed knowledge of memory representations in the human MTL, where cells are known to represent aspects of declarative memories such as the familiarity and the identity of a stimulus ^9,35,36^. This gap in knowledge motivated our study: how are MTL-based representations recruited depending on task demands through interactions with the MFC?

To address these open questions, we utilized simultaneous recordings of single neurons and local field potentials in the human MFC and MTL. We showed images to subjects and asked them to provide a “yes” or “no” answer, with either a button press or eye movement, for one of two possible tasks: a visual categorization task ^10^ or a new/old recognition memory task ^9^ (**Fig. 1a,b**). Subjects received no training and were informed explicitly of the type of task and response modality to use before the start of every block. This approach allowed us to isolate the effects of explicit task demands from sensory stimuli and motor response type. Based on a combination of single-neuron, population, and spike-field coherence analysis, we found that task demands reshape the representational geometry formed by MFC but not MTL neurons, that persistently active MFC context cells encode the currently active task, and that memory retrieval is accompanied by phase-locking between local field potentials in the MTL and spikes of memory-choice cells in MFC.

**Figure 1.**
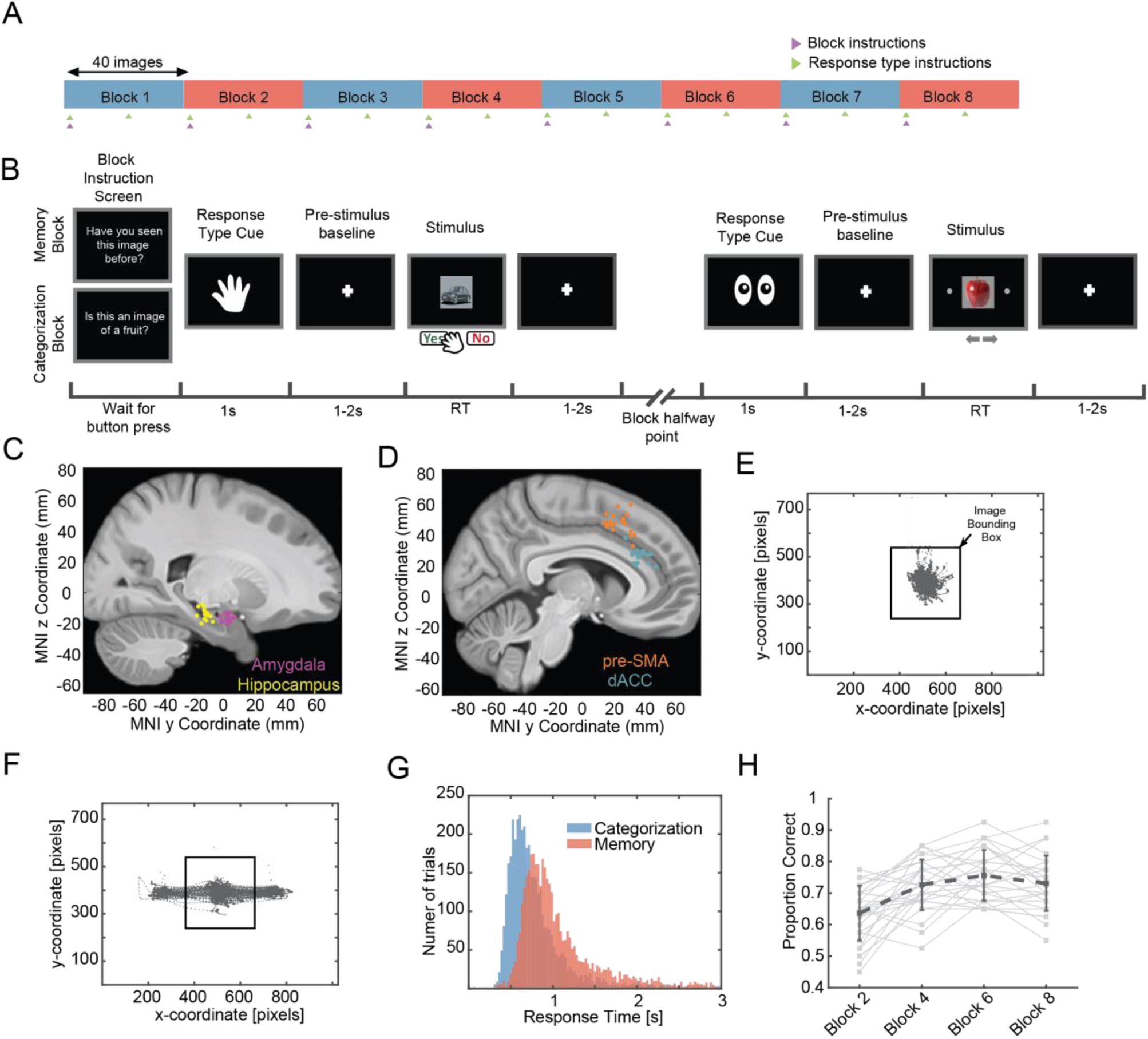
Task, electrode locations, and behavior. **(a)** Task structure. A session consisted of eight blocks of 40 trials each. The task switched with each block (blue=categorization, red=memory), and the response modality switched halfway within each block (saccade or button press; randomly assigned at the beginning of the block). The subject was explicitly instructed about the task at the beginning of each block (magenta arrow) and how to respond at the beginning and halfway point of each block (green arrow). **(b)** Example of screens shown to subject. Two example trials are shown. The fixation cross epoch between stimuli corresponds to the baseline epoch we used in some of our analyses. **(c-d)** Electrode locations in the medial temporal lobe (c) and medial frontal cortex (d). Each dot is the location of a microwire bundle in one subject. **(e-f)** Example eye tracking data from one session from the button press (e) and eye movement (f) trials. **(g)** Reaction times as a function of task pooled across all sessions (memory condition, μ ± sem, 1.27±0.02s; category condition, 0.90 ± 0.02s; p =7.6e-228, 2-sample KS test). **(h)** Memory performance improves over the course of the experiment (β = 0.56, p=8.42e-130, logistic mixed effects model). See also **Fig. S1** for an extended summary of the behavior.

## Results

### Task and behavior

We recorded from 1430 single neurons across four brain areas (**Fig. 1C-D**; see **Table S1**; 33 sessions in 13 subjects): n=203, 460, 329, and 438 neurons from anterior hippocampus, amygdala, dorsal anterior cingulate cortex (dACC) and pre-supplementary motor area (pre-SMA), respectively. We refer to hippocampus and amygdala combined as MTL (n=663 cells), and to dACC and pre-SMA combined as MFC (n=767 cells).

Subjects viewed a sequence of 320 images, grouped into 8 blocks of 40 images each, in each session (**Fig. 1A-B**). Subjects were instructed at the beginning of each block which decision to make and which response modality to use to communicate their decision. Subjects made a “yes” or “no” decision for each trial to either indicate whether an image belonged to a given visual category (“Categorization Task”) or whether an image had been seen before in the task or not (“Memory task”); no feedback was provided (see **Fig. 1A** legend and **Methods** for details on the task). Each image shown belonged to one of four visual categories: human faces, monkey faces, fruits, or cars. In each block, half of the images shown were repeated and half were novel (except in the first block, in which all images were novel). For the categorization task, the target category (“yes” response) was chosen randomly to be either human faces, monkey faces, fruits, or cars.

Choices were indicated either using saccades (leftward or rightward saccade) or button press with central fixation (**Fig. 1E-F**; mean ± s.d., 94 ± 15% of all gaze positions fell within the image shown). As expected, reaction times (RT) were significantly longer in the memory compared to the categorization task (**Fig. 1G**, mean RT of 1.48s ± 1.1 vs. 1.19s ± 1.2 respectively, p<1e-20, 2-sample KS test, mean ± s.d. across all trials in a given task). Subjects performed with an average accuracy of 97 ± 6% vs. 71 ± 6% in the categorization and memory task, respectively (mean ± s.d. across n = 33 sessions). This difference in accuracy remained after we matched for RT between the two tasks (96 ± 6% vs. 72 ± 8% with matched RT of 1.23s±0.60 vs. 1.24s±0.60 for the categorization and memory task, respectively; see methods for matching procedure; also, even without RT matching, the initial response in terms of arousal was not different between tasks as assessed by pupillometry, see **Fig. S1J-L)**. In the memory task, accuracy increased as a function of how many times an image had been shown (**Fig. 1H**, β_appearances_ = 0.56, p <1e-20, mixed effects logistic regression; also see **Fig. S1C-D** for effect of target vs. non-target on memory performance). Lastly, subjects had shorter RTs on “yes” (seen before) decisions than on “no” (novel stimulus) decisions in the memory task (**Fig. S1A**, see legend for statistics). Together, the behavior of our subjects is as expected from an MTL-dependent recognition memory task ^9^.

### Effects of task type and response modality in the medial frontal cortex

Instructions about the type of task (memory or categorization) and response modality (button press or saccade) were shown at the beginning of each block (**Fig. 1A, B**). Cells showed significant modulation of their firing rate during the baseline period as a function of task type (**Fig. 2A-B** shows an example in pre-SMA). At both the single-neuron and population level, significantly more task-type information was available in the MFC compared to MTL (25% of MFC cells vs. 12% of MTL cells, 165/767 vs. 79/663, χ^2^-test of proportions, p = 1.5e-6; **Fig. 2C**, population decoding accuracy of 98% vs. 69% in MFC and MTL, respectively, p<1e-5, Δ_true_ = 29% vs. empirical null distribution, see **Methods**). Similarly, more information about response modality was encoded by MFC vs. MTL cells (**Fig. S2E** shows an example; 14% vs 10% of cells; 84/593 vs. 59/586 in MFC and MTL respectively; χ^2^-test of proportions, p = 0.03; see; population decoding performance 84% vs. 65%, **Fig. 2D;** p<1e-5, Δ_true_ = 19% vs. empirical null distribution). Together, these results show that contextual set information (about task or response modality) is represented in the MFC during the inter-trial baseline periods.

**Figure 2.**
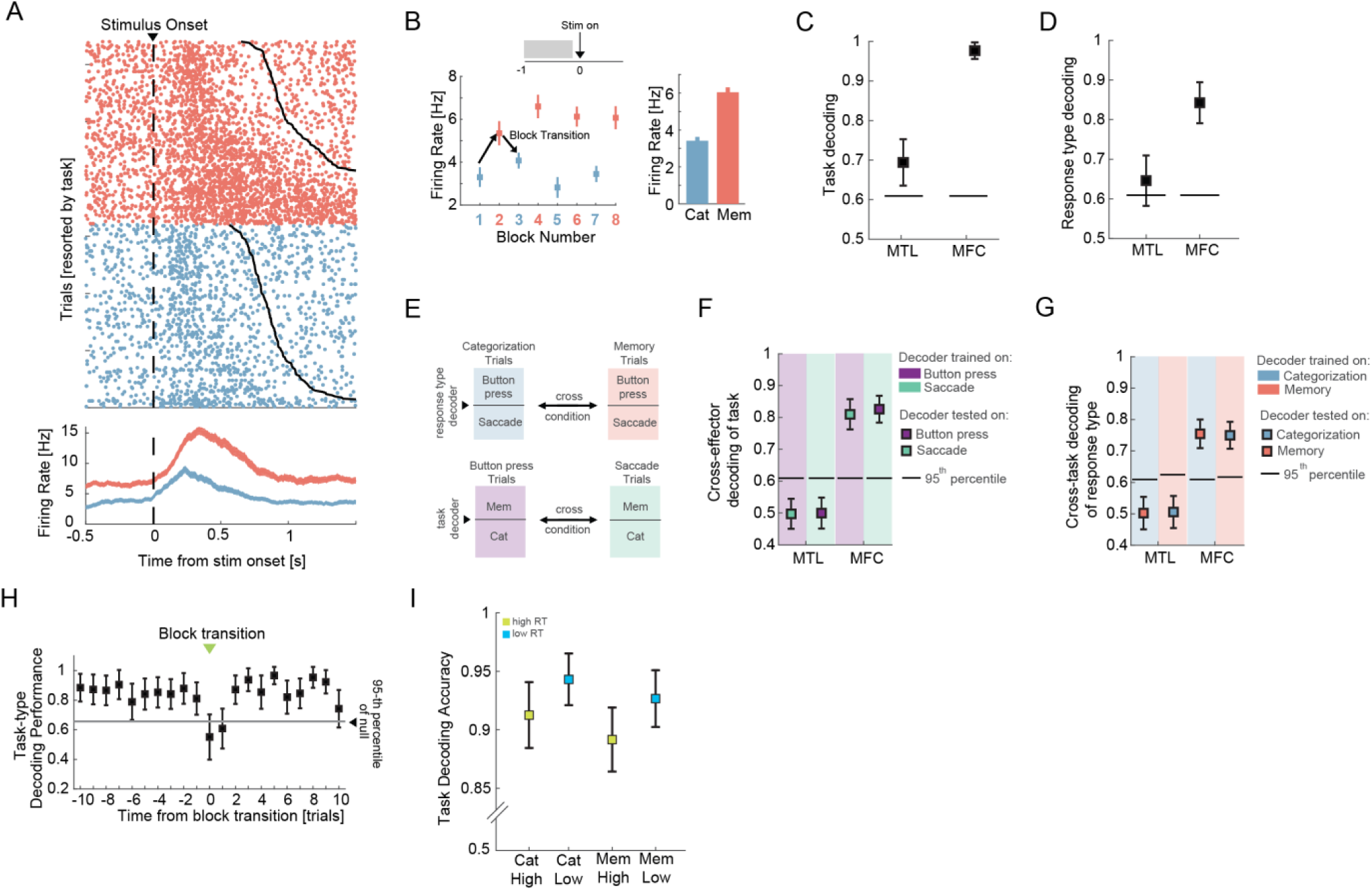
Representations of task type and response modality. **(a)** Example pre-SMA. **(b)** Average firing rate during the baseline period (−1 to 0s relative to stimulus onset) for each of the eight blocks for the cell shown in (a). The bar plot shows the average baseline firing rate, collapsed across all blocks of the same type. **(c)** Population-level decoding of task in MTL and MFC. The black lines denote the 95^th^ percentile of the null distribution (computed by shuffling the condition labels). **(d)** Population-level decoding of response modality from the baseline activity of the MFC and MTL population. Much like task type decoding, response modality was more readily decoded from the MFC population than the MTL population. **(e)** Schema of the cross-condition decoding approach. Decoding of response modality during baseline was computed across task type (memory/categorization). Similarly, decoding of task type was computed across response modality. The background color denotes the type of trials that were used to train a given decoder. For example, in the top left case, we trained a decoder to classify button press vs. saccade trials using the categorization trials (blue). We then tested the accuracy of this decoder on the memory trials (red). **(f)** Cross-response modality decoding of task type from the baseline firing rate of all recorded cells. The background color denotes which trials were used to train the decoder. The color of the tick mark denotes which trials were used to test the decoder. **(g)** Cross-task decoding of response modality. **(h)** Decoding accuracy was lower for the first two trials after a task type context switch. Shown is decoding performance as a function of trial number relative to a task type switch (green arrows in Fig. 1a; transitions from categorization to memory and vice-versa were pooled). Error bars indicate standard deviation in all panels, with the exception of panel B where they indicate the standard error of the mean. (**i**) Task decoding on trials with short reaction times was more accurate than decoding on long reaction time trials. Shown separately for categorization and memory trials (p = 2e-11 and 7e-13 respectively, Wilcoxon rank sum test). Error bars in this case denote standard error in decoding accuracy across trials (80 trials in each of the 4 groups). See **Fig. S2** for additional analyses that break down context effects by specific anatomical regions.

After a task switch, contextual signals emerged rapidly within 1-2 trials in the new context (**Fig. 2H**). Task switching costs are also reflected in the subjects’ longer reaction times shortly after a change in task or effector type (**Fig. S2A**). Furthermore, task type representations during the baseline period were stronger on trials where the subject produced a fast response versus those where the response time was slow (**Fig. 1I**), indicating behavioral relevance. Lastly, we tested whether the two types of contextual signals were sufficiently robust to avoid interference with one another, using a cross-condition generalization decoding analysis ^37^ (**Fig. 2E**). We first trained a decoder to discriminate task type on trials where the subject was instructed to reply with a button press, and then tested the performance of this decoder on trials where the subject was instructed to use saccades (and vice-versa). This analysis revealed that the two neural representations generalize in the MFC but not in the MTL (**Fig. 2F-G)**. Together, this shows that the contextual signals in MFC were robust and behaviorally relevant.

### Cross-condition generalization of memory and image category

We next asked whether the neural representations of image category and familiarity are sensitive to task demands. At the single-unit level, we examined visually-selective (VS) cells ^35^, whose responses are thought to reflect input from high level visual cortex, and memory-selective (MS) cells ^9^, whose response signals stimulus familiarity (**Fig. 3A-B** shows examples). Of the MTL cells, 40% were visually selective (264/663) and 11% (73/663) were memory selective (see Methods for selection models). By comparison, in the MFC, 13% (103/767) of the cells were visually selective and 11% (84/767) were memory selective. We first performed single-neuron analysis of the selected MTL cells and found that both visual and memory selectivity was present in both the memory and categorization blocks (**Fig. S3D, E** and **S3G-I**). We next took a population-level approach (over all single units, without selection) to answer the question whether the same or different groups of cells represent visual- and memory information across tasks (**Fig. 3C-H**). We used decoding accuracy based on condition averages to quantify the extent to which a particular type of information was represented (see Methods; single-trial decoding reveals quantitatively similar results, see **Fig. S4D-E**). In both MTL and MFC, image category could be decoded well above chance (**Fig. 3C**, 100% and 58% in MTL and MFC respectively, chance level = 25%). In the MTL, the ability to decode category was not affected by the task (**Fig. 3C**, 100% for both tasks). In contrast, in the MFC, decoding accuracy for image category was significantly higher in the memory task (**Fig. 3C**, 72% vs. 43%). Similarly, memory (i.e. new vs. old) was decodable in both the MTL and MFC (**Fig. 3D**, 80% vs. 84% respectively, chance level = 50%), with no significant difference in decoding accuracy between the two tasks in MTL (**Fig. 3D**, 79% vs. 80% in categorization and memory trials respectively) and significantly better decoding ability in MFC in the memory task (**Fig. 3D**, 96% vs. 73%).

**Figure 3.**
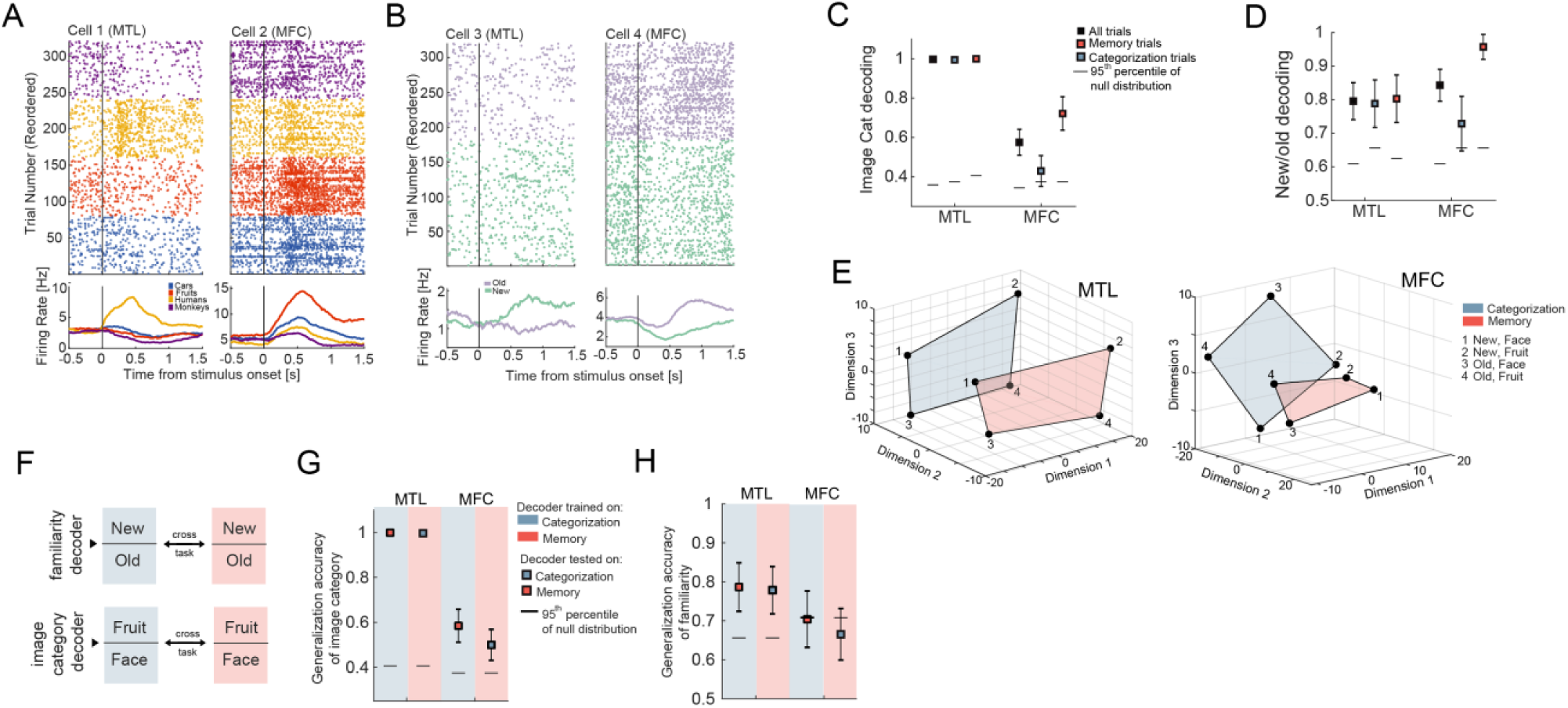
Task-independent representations of image category and familiarity (new/old). **(a-b)** Example cells. **(a)** Cells whose activity represents image category. **(b)** Cells whose activity differentiates between new and old stimuli. **(c)** Decoding accuracy of image category from all recorded cells was significantly higher in the MTL (left) relative to MFC (right). Mean decoding accuracy ± std is shown for all trials (black tick mark), categorization trials (blue tick mark) and memory trials (red tick mark). Black lines show 95^th^ percentile of the null distribution, constructed by shuffling the image labels. **(d)** Decoding of new vs. old was possible at similar accuracy in both MTL and MFC. Decoding accuracy is shown for all trials and separately for categorization and memory trials (using ground truth labels for all trials). Black lines show the 95^th^ percentile of the null distribution. **(e)** Population activity of all MTL (left panel) and MFC (right panel) cells, collapsed to three dimensions using MDS (Euclidean distance). Individual points show the mean activity of the population for that specific condition. The highlighted plane contains all locations of state space occupied by a given task for the case of fruits vs. faces as the binary category distinction (for illustration only; all analysis uses all categories). Note how the geometry of the representation allows for a decoder that is trained on one task to generalize to the other task (see **Fig. S4C** for example decoder hyperplanes in this geometry). **(f)** Schematic of the approach used for the cross-condition generalization analysis. Color indicates task (blue=categorization, red=memory). (Top) We trained a decoder to discriminate between new vs. old trials on categorization trials and then tested its performance on new vs. old stimuli encountered during the memory condition (and vice-versa). (Bottom) Similarly, a decoder that is trained to discriminate between image categories (in this example face vs. fruits, all results include all 6 possible pairs) on categorization trials, was tested on memory trials. **(g)** Cross-condition generalization performance for image category decoders indicates that category coding is independent of task type. The background color (blue or red) indicates the condition in which the decoder was trained. The color of the tick mark indicates the task on which the decoder was tested. The black lines indicated the 95^th^ percentile of the null distribution, constructed by shuffling the image category labels. **(h)** Cross-condition generalization performance for decoders trained on new vs. old classification indicates task-independent encoding of familiarity only in MTL, but not MFC.

These results indicate that information about familiarity and stimulus category in the MTL is independent of task demands, whereas it is sensitive to task demands in the MFC. To gain insight into the geometry of the population-level representations, we next assessed whether the decoders trained to report familiarity and the category of the stimuli in one task would generalize to the other task. This would indicate that familiarity and category are represented in an abstract format ^37^ (**Fig. 3F**). First, cross-task generalization performance was greater in the MTL than MFC for both image category (**Fig. 3G**; 100% vs. 54%, averaged across the two cross-condition decoding performances) and familiarity (**Fig. 3H**; 79% vs. 67%). This suggests that the degree of generalization of these two variables is higher in the MTL than in the MFC. Note that in the MTL, familiarity decoding was supported by amygdala neurons (**Fig. S4A-B**). Second, to help understand the geometry of these neural representations, we projected the average MTL and MFC population activity for all possible pairings of familiarity, image category, and task (8 different conditions) into a 3D state-space using multi-dimensional scaling. For illustration purposes, we show this for the two image categories (fruits, faces) for which memory performance was the best (for all other analyses, we used all image categories). This revealed that in the MTL (**Fig. 3E**, left), the relative positions of a “new face” with respect to an “old face” is preserved across tasks (shown as different colored planes). The relatively parallel location of the subspace of neural activity occupied by the two tasks permits across-task generalization. This geometry is similar to the one observed in monkeys in a different task and discussed in ^37^. In contrast, in the MFC (**Fig. 3E**, right), the relative positions of the four conditions are not preserved, which is consistent with the context sensitivity in MFC but not in MTL (**Fig. 3G-H**), suggesting a reason for the cross-task generalization performance. Together, these analyses show that the representation of image category and memory is the same across tasks in MTL, whereas in MFC representations were sensitive to task demands, a phenomenon that we investigated next in more detail.

### Representation of choice in the MFC

We next investigated how the subject’s choice (yes or no) is represented. Cells in the MFC show modulation of activity with the subject’s choice (**Fig. 4A** shows examples). This signal was strongest in the MFC, with average population decoding performance of 89%, compared to 68% in the MTL (**Fig. 4B**, “yes” vs. “no” decoding). Choice decoding was strongest shortly after stimulus onset well before the response was made (**Fig. S7A)**. To disassociate representation of choice (yes vs. no) from the representation of ground truth (old vs. new) during the memory recognition task, we fit a choice decoder to a set of trials that were half-correct and half-incorrect. We found that the activity of MFC cells predicted choice, but not the ground truth (**Fig. 4C**). A similar analysis revealed that choice can be decoded separately from both correct and incorrect trials (**Fig. S5I**). As a control for potential confounds due to RT differences between tasks (see **Fig. S1A**), we acquired data from a separate control task in which we eliminated RT differences behaviorally by adding a waiting period (6 sessions in 5 subjects; n=180 and 162 neurons in MTL and MFC, respectively; see **Methods** and **Fig. S6**). Like in the original task, MFC cells represented the subject’s choice (**Fig. S6C-G**), thereby confirming that this separation is not due to RT differences.

**Figure 4.**
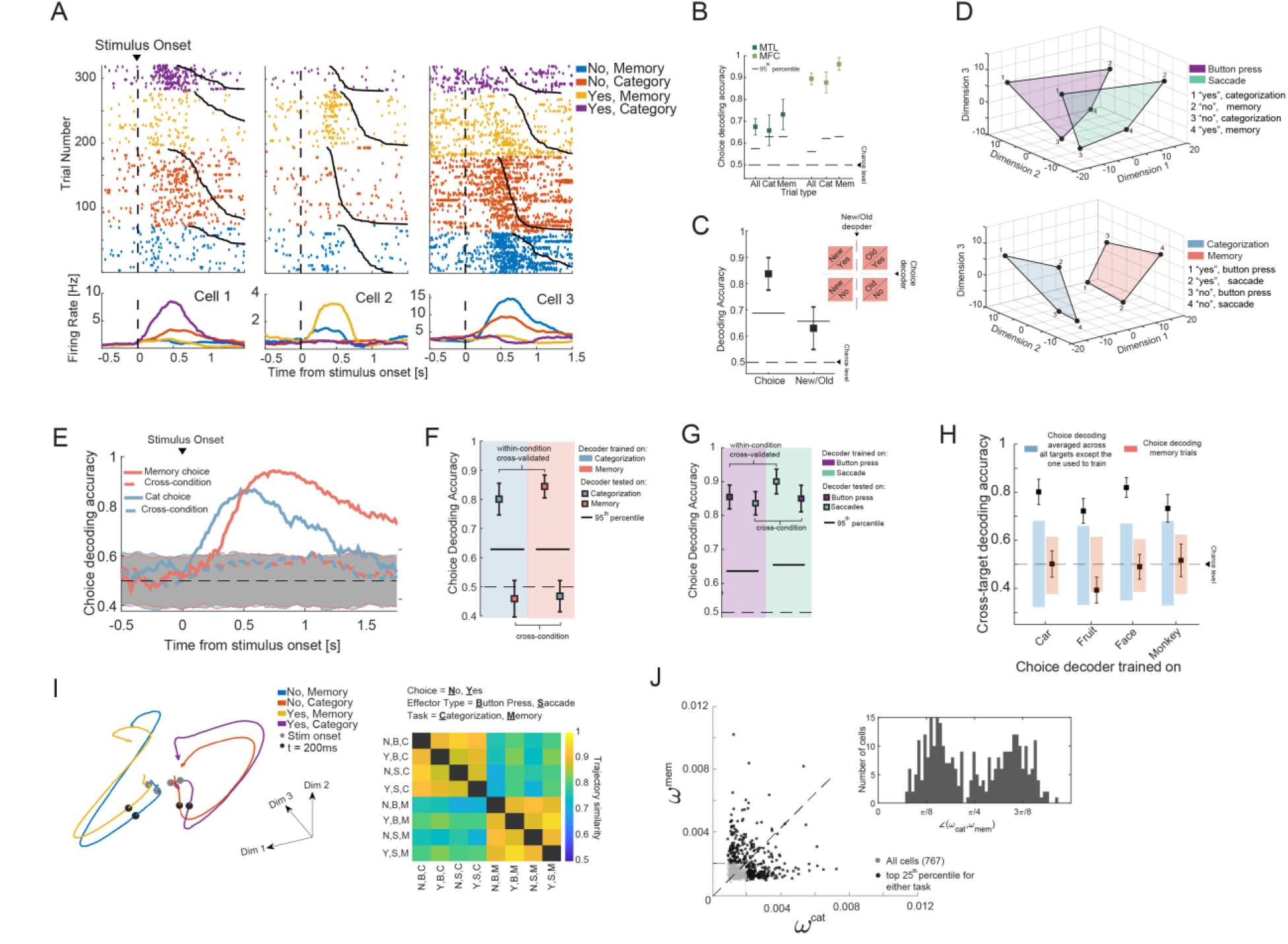
Task-specific representation of choice in MFC. **(a)** Responses of choice cells in the MFC are diverse. Shown are three example cells, split by choice (yes or no) and task. **(b)** Population choice decoding accuracy was significantly greater in MFC compared to MTL. **(c)** MFC cells represent the choice during the memory trials and not the ground truth (new/old). We sampled an equal number of memory-task trials with the following label combinations of familiarity and choice: (1) new/no (correct), new/yes (incorrect), old/no (incorrect), old/yes (correct). Notice that in half the trials, familiarity and choice are congruent and in the other half they are incongruent. We fit separate decoders to read-out choice and new/old from the population. Choice can be decoded well above chance levels, whereas new/old cannot. **(d)** Population summary (neural state space) of choice-related activity in MFC, plotted in 3D space derived from MDS. Each point indicates the mean activity of a choice in one of the two tasks. (Top) Variability due to response modality. The highlighted planes connect the points of state space occupied by activity when utilizing button presses (purple) or saccades (green). The relative positions of the points stays the same, suggesting generalization of choice encoding across response modality. (Bottom) Variability due to task type. The highlighted plans connect the points of state space occupied by activity in the same task (red = memory, blue = categorization). The relative position of the 4 points is not preserved across the task planes, suggesting lack of generalization across task. **(e)** Choice-decoders trained in one task do not generalize to the other task. Shown here as a function of time (500ms bin, 16ms step size). See **Fig. S7** for results individually for dACC and pre-SMA. **(f)** Same as (e) but shown for a fixed time window [0.2 1.2] seconds relative to stimulus onset. As suggested by panel d (bottom), decoding does not generalize across tasks. To avoid reaction time confounds, this analysis is shown only for cells recorded in the control task, in which RT was equated due to a go cue (see methods and **Fig. S6**). **(g)** Decoding generalizes across effectors as suggested by panel d (top). Here, performance is shown averaged across categorization and memory choices (chance level = 50%). As in (f), decoders are fit using cells recorded during the control task. **(h)** Generalization between different sub-tasks of the categorization task but not between task types. Average yes vs. no decoding performance for decoders trained in subsets of categorization trials (x-axis). The blue and red shading shows the range of the null distribution estimated by shuffling the labels. **(i)** (Left) State-space trajectories for the four conditions arising from the combination of response (yes, no) and task (categorization, memory). (Right) Trajectory similarity, computed in a 8D latent space (recovered using GPFA, see Methods) across the eight conditions arising from the combinations of choice, effector type, and task. The similarity is averaged in the time window [0 500ms] after stimulus onset. Latent factors were recovered from the condition averages, across all MFC cells. **(j)** Scatter plot of the decoder weight assigned to each cell in decoding choice during the categorization task (x-axis) and memory choice (y-axis). The cells in the top 25-th percentile are shown in black. The inset shows the angle created by the vector 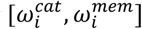 with respect to the x-axis of the cells marked in black.

We used multi-dimensional scaling to visualize the population activity for the eight combinations of choices, task types, and response modality (**Fig. 4D**, see **Methods**). The resulting geometrical configuration indicates that choice decoding generalizes across response modality (**Fig. 4D**, top) but not across task types (**Fig. 4D**, bottom). To test this hypothesis, we computed the cross-task generalization performance of a decoder trained on choices during one task and tested on the other. We performed this analysis across time (**Fig. 4E)**, as well as in a single post-stimulus time bin (**Fig. 4F)**. To avoid confounds due to response time differences, we performed the fixed window analysis (**Fig. 4F**) only for the control task, where the timing between tasks was identical (**Fig. S6B**). Importantly, while the choice signal does not generalize across task types, it does generalize across response modality within the same task type (**Fig. 4G**). To test the possibility that any task might yield a unique choice axis that does not generalize to any other task, we considered the four subtasks that make up the categorization trials in a given session (the target category can be any one of the four possible image categories). We tested whether the choice signal generalizes across these subtasks by training and testing across blocks requiring different categorizations. We found that choice decoding generalized across all sub-tasks in the categorization task, but not the memory task (**Fig. 4H)**. Next we compared the dynamics of the population activity between the eight conditions arising from the combination of choice, effector type, and task in a state-space model recovered using Gaussian Process Factor Analysis (GPFA ^38^, see **Methods**). A comparison of the pairwise similarity between the trajectories in state space (**Fig. 4I**, left) in the first 500ms after the stimulus onset reveals that the patterns of dynamics in state space cluster first by task type (**Fig. 4I**, right; also see **Supplementary Video 1**).

We next examined whether the population-level analysis relied on different sets of neurons to decode choice in each of the two tasks. We did this by determining how individual cells are recruited by a linear decoder ^37,39^. For each cell, we quantified its importance ^39^ (see methods) for both the memory and categorization choice decoder (**Fig. 4J)**. We then plotted the degree of specialization for each cell based on its importance in each task (see **Methods**). Cells that report choice independently of task should lie on the diagonal (i.e. an angle of π/4). Instead, we found that the distribution of angles is significantly bimodal across all cells (**Fig. 4I**, inset plot, p<1e-5, Hartigan dip test), with modes centered away from the diagonal. Despite this bimodality, we could still use the cells that are the most *useful* in one task, to train a new decoder that can predict choice well above chance (although significantly weaker) in the other task (**Fig. S7C**). Note that this is not an example of cross-task generalization, since we are fitting a new decoder.

Taken together, these results reveal a task-dependent and highly flexible utilization of different subspaces of neural activity that represents choices in the MFC.

### Task-dependent spike-field coherence between MFC cells and MTL LFP

It is thought that the selective routing of decision-related information between MTL and MFC is coordinated by inter-areal spike-field coherence ^6^. We therefore next asked whether MFC neurons phase-lock to ongoing oscillations in the LFP in the MTL and, if so, whether the strength of such interactions is modulated by task demands. We performed this analysis for the 13 subjects and 33 sessions for which we simultaneously recorded from both areas (**Fig. 5A** summarizes our approach; see methods for details on preprocessing of the data). In the following, we only utilized neural activity from the 1s baseline period that precedes stimulus-onset to avoid confounds related to stimulus-onset evoked activity. Individual cells in the MFC showed strong task modulation of MFC to MTL spike-field coherence (**Fig. 5B** shows a single-cell example in the dACC). At the population level, MFC cells showed significantly stronger theta-band coherence with MTL oscillations during the memory compared to the categorization task (**Fig. 5C**, 8822 cell electrode pairs; p = 1.3e-7, paired t-test, measured at 5.5Hz). This was the case for both MFC-hippocampus, and MFC-amygdala interactions (**Fig. 5D**, n = 3939, p = 8.8e-4; n = 4884, p = 4.3e-5 respectively, paired t-test). This effect was due to changes in phase preference as there was no significant difference in MTL LFP power between the tasks (**Fig. 5E**, p = 0.08, paired t-test of signal power at 5.5Hz, estimate across all 8822 cell-electrode pairs). Out of the 767 MFC cells, a significant subset of ∼100 cells were phase-locked to the theta-band MTL LFP (**Fig. S9A**), with the largest proportion preferring 3-8Hz.

**Figure 5.**
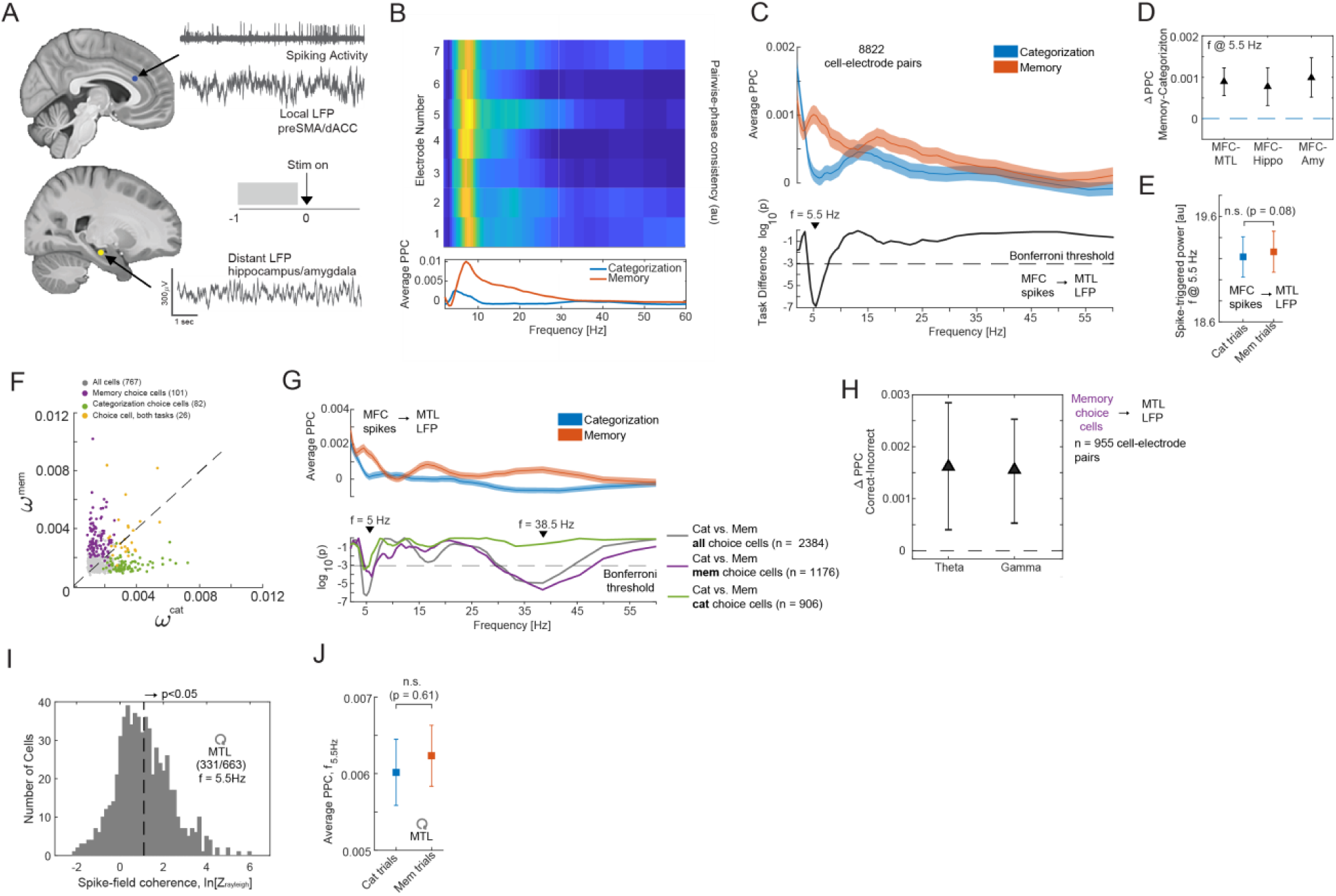
Coherence between MFC spikes and MTL field potentials is modulated by task demands. **(a)** Schematic of the spike-field coherence analysis. (top) Example of LFP and spiking activity from the dorsal anterior cingulate cortex. (bottom) Example LFP recording from the hippocampus. Inset illustrates that throughout this figure, only the spikes from the 1s long baseline period before stimulus onset were used for analysis. **(b)** Example spike-field coherence for a dACC cell with all recorded channels in the ipsilateral hippocampus. Bottom shows average across all pairs separately for memory and categorization trials. **(c)** MFC cells increase their phase locking to MTL LFP during the memory relative to the category task. (top) Average inter-area pairwise phase consistency (PPC) of all cell-electrode pairs, split by task (red = memory, blue = categorization). (bottom) p-value as a function of frequency, testing differences between the two tasks. The peak difference between the two task conditions occurs as f = 5.5Hz. Dashed line shows the threshold we used (0.05/56 Bonferroni corrected). **(d)** Difference in average inter-area PPC, measured at 5.5Hz, between the two task conditions, shown for all MFC and MTL cell-electrode pairs (n = 8822; paired t-test, mem vs. cat, p = 1.3e-7), MFC-hippo (n = 3938; p = 8.8e-4), and MFC-amygdala (n = 4884; p = 4.3e-5). The error bars denote the 95% confidence interval of the mean difference in PPC between the two tasks, as estimated via bootstrapping (n = 10000 iterations). **(e)** Average spike-triggered power for all cell-electrode pairs, shown separately for categorization- and memory trials. There was no significant difference in the spike-triggered power in the two task conditions (paired t-test, n = 8822 cell electrode pairs, p = 0.08). Error bars indicate the s.e.m. **(f)** Single-neuron analysis. Scatter plot of the normalized weight (importance index, see methods) assigned by the decoder to each cell when decoding choices in either the memory or categorization task. Memory and categorization choice cells are indicated in color (see Methods for the selection model that was used to identify these cells). **(g)** MFC-MTL spike-field coherence for choice cells. (top) Average PPC for all choice cells in MFC (209 cells, 2384 cell-electrode pairs) for memory-(red trace) and categorization trials (blue trace). (bottom) Significance of PPC differences between the memory and categorization task as a function of frequency, shown separately for memory choice cells (purple trace; 1176 cell-electrode pairs) and categorization choice cells (green trace; 906 cell-electrode pairs). Both memory and categorization choice cells show a strong theta-band difference in coherence between the two tasks, but only the memory choice cells show a difference in the gamma band (−6e-04 vs. 0.001 in categorization and recognition trials respectively, p = 2e-6, t-test). **(h)** Baseline PPC difference between correct and incorrect trials during the memory task shown for the theta (f = 5Hz; p = 0.009, paired t-test) and gamma band (f = 38.5Hz; p = 0.002, paired t-test). The difference is computed for memory choice cells only (purple cluster from **Fig. 5F**). The error bars denote the 95% confidence interval of the mean difference in PPC as estimated via bootstrapping (n = 10000 iterations). **(i)** Spike times of MTL cells are coherent with local theta-band (3-8 Hz, centered at 5.5 Hz) LFP recorded in the same area in which a cell was recorded from. Shown is a histogram of the log of the Rayleigh z-value for all recorded MTL cells. We computed the spike-field coherence in the theta range (3-8Hz, center at 5.5 Hz; phase determined from extracted spike snippets) using all spikes of a cell during the entire experiment (see methods). **(j)** Average local PPC in the MTL did not differ as a function of the task in the MTL (f = 5.5Hz; p = 0.51, paired t-test, n = 2321 cell-electrode pairs).

To determine if there is a relationship between the tuning of cells in MFC and their inter-area coherence with MTL, we selected for choice cells independently in the categorization and memory task (see **Methods** for selection model; note selection controls for RT differences). This revealed that 101/767 and 82/767 cells are significantly modulated by choice during the memory and categorization task, respectively (p<0.001 vs. chance for both; see **Fig. 4A** Cell 2 and 1 for an example, respectively). Single-unit decoding showed that choice cells selected in the memory task do not carry information about the subject’s choice in the categorization task and vice-versa (**Fig. S5A-D**). In addition, the removal of the selected choice cells from a population decoding analysis with access to all recorded neurons significantly diminishes decoding performance (**Fig. S5F, G**). Importantly, each of the selected cells has a high importance index, as determined from the population decoding (**Fig. 5F)**. Considering only the MFC choice cells revealed that this subset of cells were similarly increasing their phase-locking during the memory task (**Fig. 5G, top**), with the strongest effect again in the theta range (f_peak_ = 5 Hz, p = 1e-6, paired t-test). Both categorization and memory choice cells showed this pattern of modulation (**Fig. 5G**, bottom). In addition, the memory choice cells exhibited an increase in gamma band coherence (**Fig. 5G**, f_peak_ = 38.5Hz; p = 2e-6, paired t-test). The baseline theta and gamma synchrony of memory choice cells were predictive of the accuracy of the choice made for the picture shown in that trial (**Fig. 5H**, see legend for statistics). Lastly, to exclude the possibility that this inter-area effect was due to task-dependent changes within the MTL, we examined the phase-locking properties of MTL cells to their own locally recorded LFP (LFP and spiking activity is recorded on separate electrodes, see Methods). The spiking activity of 331/663 MTL cells was significantly related to the theta-frequency band LFP (**Fig. 5I**, shown for f = 5.5 Hz). The strength of this local spike-field coherence was, however, not significantly different between the two tasks (**Fig. 5J;** p = 0.61, paired t-test, n = 2321 cell-electrode pairs). Together, this shows that MFC to MTL spike-field coherence is selectively increased during the memory relative to the categorization task, thereby showing that this mechanism of information routing can be selectively up-regulated by task demands.

## Discussion

We investigated the nature of flexible decision-making in the human brain by probing how the geometry ^37^ of neural representations of stimulus memory, stimulus category, and choice is modified when subjects switch between tasks. We found a striking difference between brain areas: In the MTL, representations of stimulus memory and stimulus category were task-demand independent, whereas in the MFC, representations of stimulus memory, stimulus category, and choices were all highly sensitive to task demands. A group of task demand-dependent cells in the MFC that we reveal here are choice cells, which signal behavioral decisions preferentially for either memory- or categorization decisions irrespective of response modality and regardless of the ground truth of the decision. At the population level, choices in both the memory and categorization task were decodable with high reliability, but these decoders did not generalize across the two tasks. Thus, from the point of view of downstream areas, neurons within MFC formed two separate decision axes: one for memory-based decisions and one for categorization-based decisions. Moreover, these two decision axes were instantiated selectively so that they were only present when the current task required them.

These findings contrast with prior work on task switching between different tasks that required purely perceptual decisions, which found a single decision axis in monkey pre-frontal cortex, with task-irrelevant attributes also represented ^40^. Here we show that memory-based choices add a second decision axis, which is present only when decisions engage memory retrieval processes. While task-sensitive representations of choice have been shown in recordings from nonhuman primates during perceptual decision-making ^10,40,41^, our data for the first time find choice representations that specifically signal declarative memory-based choices. It has long been appreciated that the frontal lobes are critical for initiating and controlling memory retrieval ^34,42,43^, and it is thought that the memory retrieval network implements a set of specialized processes separate from those utilized for other kinds of decisions ^13,44,45^. The memory-choice axis we reveal here is direct evidence for this hypothesis and reveals a potential cellular substrate for this critical but poorly understood aspect of human cognitive flexibility.

A second group of cells we characterized in MFC signal the currently relevant goal (task type and response modality) throughout the task. These cells switched their activity pattern when instructions indicated a change in task demands. While these switches were rapid, they were not instantaneous, likely reflecting the cost of switching between memory retrieval and categorization modes ^46-48^. We hypothesize that these cells facilitate holding in working memory ^49,50^ the active task set and configure brain networks in preparation for appropriate execution of the instructed task ^23,51,52^. Task switching costs are a much investigated aspect of cognitive flexibility ^8,46-48^, but it remains little understood how they arise, and why some task switches are more difficult than others. The MFC cells we describe here offer an avenue to directly investigate these critical questions.

Finally, we uncovered a possible mechanism by which memory-based information can be routed dynamically between MFC and MTL when a task requires memory retrieval. Changing long-range synchronization of neural activity is thought to be a way by which functional connectivity between brain areas can be changed flexibly ^4,53-55^. Here, we reveal a specific instance of this phenomenon at the cellular level in humans in the form of changes in the strength of cortico-MTL functional connectivity. While hippocampal-mPFC functional connectivity in rodents supports spatial working memory ^6^ and is prominent during both spatial navigation and rest ^56-58^, it remains unknown whether this pathway serves a role in long-term memory retrieval and, if so, whether it can be engaged selectively. Here, we show that MFC-MTL theta-frequency connectivity is selectively enhanced during memory retrieval. In addition, memory choice cells in MFC exhibited enhanced gamma-frequency band coordination of their spiking activity with the MTL LFP. This reveals a specific cellular-level instance of a role for gamma oscillation-mediated coordination of activity between distant brain regions ^6,59^ in human memory retrieval. Disrupted MFC-MTL functional connectivity is a key feature of impaired executive control in psychiatric disease ^60^ and is a pertinent phenotype of genetic mouse models of human schizophrenia ^29^. Our findings suggest the hypothesis that a specific impairment that might result from this impaired connectivity is an ability to selectively initiate memory retrieval when demanded by a task.

## Acknowledgments

We would like to thank the members of the Adolphs and Rutishauser labs for discussion on the task and results. Additionally, we would like to thank Columbia Theory Center members Fabio Stefanini, Mattia Rigotti, and Marcus Benna for their population decoding analysis expertise. We thank all subjects and their families for their participation. This work was supported by NIMH (R01MH110831 to U.R.), the Caltech NIMH Conte Center (P50MH094258 to R.A.), the National Science Foundation (CAREER Award BCS-1554105 to U.R.), and a Memory and Cognitive Disorders Award from the McKnight Foundation for Neuroscience (to U.R.).

## Author Contributions

J.M., U.R., and R.A. designed the study. J.M. performed the experiments. J.M., S.F, and U.R. analyzed the data. J.M., U.R., R.A., S.F. wrote the paper. A.M. performed surgery and supervised clinical work.

## Methods

### 1. Subjects

Subjects were 13 adult patients being evaluated for surgical treatment of drug-resistant epilepsy that provided informed consent and volunteered for this study (see **Table S1**). The institutional review boards of Cedars-Sinai Medical Center and the California Institute of Technology approved all protocols. We excluded potential subjects who did not have at least one depth electrode in medial frontal cortex.

### 2. Electrophysiology

We recorded bilaterally from the amygdala, hippocampus, dACC, and preSMA using microwires embedded in hybrid depth electrodes^61^. From each micro-wire, we recorded the broadband 0.1-9000Hz continuous extracellular signals with a sampling rate of 32 kHz (ATLAS system, Neuralynx Inc.). Subjects from which not at least one well identified single-neuron could be recorded were excluded.

### 3. Spike sorting and single-neuron analysis

The raw signal was filtered with a zero-phase lag filter in the 300-3000Hz band and spikes were detected and sorted using a semi-automated template-matching algorithm ^62,63^. All PSTH diagrams were computed using a 500ms window with a step-size of 7.8ms. No smoothing was applied.

### 4. Electrode localization (relevant for Figure 1)

Electrode localization was performed based on post-operative MRI scans. These scans were registered to pre-operative MRI scans using Freesurfer’s mri_robust_register ^64^ to allow accurate and subject-specific localization. To summarize electrode positions and to provide across-study comparability we in addition also aligned the pre-operative scan to the MNI152-aligned CIT168 template brain ^65^ using a concatenation of an affine transformation followed by a symmetric image normalization (SyN) diffeomorphic transform ^66^. This procedure provided the MNI coordinates that are reported here for every recording location.

### 5. Eye tracking (relevant for Figure S1)

Gaze position was monitored using an infrared-based eye tracker with a 500Hz-sampling rate (EyeLink 1000, SR Research)^67^. Calibration was performed using the built-in 9-point calibration grid and was only used if validation resulted in a measurement error of <1 dva (average validation error was 0.7 dva). We used the default values for the thresholds in the Eyelink system that determine fixation and saccade onsets.

### 6. Task

Each session consisted of 8 blocks of 40 trials shown in randomized order. At the beginning of each block, an instruction screen told subjects verbally the task to be performed for the following 40 trials (categorization or recognition memory), the response modality to use (button presses or eye movements), and which visual category is the target (for categorization task only; either human faces, monkey faces, fruits, or cars; order was pseudo-random so that each image type was selected as the target at least once) (see Figure 1). The task to solve was either “Have you seen this image before, yes or no?” or “Does this image belong to the target category, yes or no”. Odd-numbered blocks (1,3,5,7) were categorization blocks; even numbered blocks were memory blocks (2,4,6,8). Button presses (yes or no) were recorded using a response box (RB-844, Cedrus Inc.). Eye movements to the left or right of the image served as responses in the eye movement modality (left=yes, right=no). The mapping between button and screen side and yes/no responses was fixed and did not change; “yes” was on the left and “no” was on the right. Subjects were reminded that left = yes, and right = no, at the beginning of each of the 8 blocks. In the first block, all images were novel (40 unique images). In all subsequent blocks, 20 new novel images were shown randomly intermixed with 20 repeated images (the “old set”). The 20 repeated images remained the same throughout a session. We used entirely non-overlapping image sets for patients that completed multiple sessions. The response modality (button presses or eye movements) was selected randomly initially and switched in the middle of each block (an instruction screen in the middle of each block showed the response modality to be used for the remainder of the block). In sessions where eye tracking was not possible due to problems with calibration (5 sessions in 3 patients; see **Table S1**), all trials used the button presses as the response modality. No trial-by-trial feedback was given. In between image presentations, subjects were instructed to look at the fixation cross in the center of the screen.

### 7. Control Task (relevant for Figure S6)

In 5 of the 13 subjects in this dataset (6/33 sessions), we ran an additional control task in order to help determine if neural responses reflected processing of stimuli, of decision variables, or of motor response plans. Unlike the standard task where the subjects could respond at any time after the stimulus onset (thus making it difficult to distinguish decision from choice), in this control task the subjects were instructed to wait until the response cue in order to register their answer, either with a button press or with a saccade. The stimulus was presented for a fixed amount of time (1s duration) and after a 0.5 – 1.5s delay period, the subjects were asked to respond to the question relevant for that block.

### 8. Mixed-effects modeling of behavior (relevant for Figures 1, S1)

For the group analysis of behavior, we used mixed-effects models of the form *y* = *Xβ* + *Zb* + *ε*, where y is the response, X is the fixed-effects design matrix, *β* is the fixed-effects coefficients, Z is the random-effects design matrix, b is the random-effects coefficients, and *ε* is the error vector. In all analysis, we used a random intercept model with a fixed slope. The grouping variable for the random-effects was the session ID. The reported p-values in the main text correspond to the fixed-intercept for the relevant variable. In the case of measuring the effect of number of expositions (i.e. number of times an image was seen) on the subject’s accuracy during the memory trials, we used a mixed-effects logistic regression with the independent variable as an ordinal-valued whole number ranging from 1-7. The response was a logical value indicating success or failure on each memory question. Prior to running any analysis of reaction time data, we excluded outliers from the distribution using the following procedure: a sample was considered an outlier if it was outside the 99^th^ percentile of the empirical distribution.

### 9. Reaction time matching procedure

As a control, we matched for RTs between the two tasks (categorization and memory) to exclude for potential differences due to difficulty. To achieve this, we first added noise to all reaction times (s.d. = 1ms), followed by locating pairs of trials with RTs that were equal to within a tolerance of 0.1s. Matching pairs were then removed and this procedure was repeated iteratively until no further matches could be found. Unmatched trials were excluded (resulting in reduced statistical power due to fewer trials available). We only used the resulting match if the RTs between the two groups were not significantly different. If not, the procedure above was repeated.

### 10. Selection of visually (VS) and memory-selective (MS) cells (relevant for Figure S3)

A cell was considered as a VS cell if it response co-varied significantly with visual category as assessed using a 1×4 ANOVA test at p<0.05. For each selected cell, the preferred image category was set to be the image category for which the mean firing rate of the cell was the greatest. All trials were used for this analysis. MS cells were selected using the following linear model:

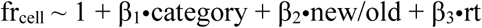

where *category* is a categorical (1×4) variable, *new/old* is a binary variable, and *rt* is a continuously valued variable. A cell was determined to be memory selective if the t-statistic for β2 was significant with p<0.05. We excluded the first block of trials (40 images) from the analysis, in order to keep the number of new and old stimuli the same. Spikes were counted for every trial in a 1s window starting at 200ms after stimulus onset.

### 11. Selection of choice cells (relevant for Figure S5)

Choice cells were selected using a regression model applied to the firing rate in a 1s size window starting 200ms after stimulus onset. We fit the following regression model:

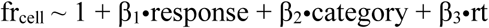

where the response is binary (yes or no), category is a categorical variable with four levels, and RT is the reaction time. We fit this model separately to trials in the memory-and categorization condition, assuring independent selection of cells. RT was included as a nuisance regressor to control for reaction time differences between the two possible responses (see **Fig. S1a)**. A cell qualified as a choice cell if the t-statistic of the β1 term was significant at p<0.05 for at least one of the two task conditions. The response preference of significant cells for either yes or no was determined based on the sign of β1 (positive = yes, negative = no). Notice that the selection process uses separate trials for memory choice cells and categorization choice cells. All trials regardless of whether the answer was correct or incorrect were used for selection. To estimate the significance of the number of selected cells, we generate a null distribution by repeating above selection process 1000 times after randomly re-shuffling the response label. We estimated this null distribution separately for choice cells in for the memory-and categorization condition and used each to estimate the significance of the number of selected cells of each type.

### 12. Chance levels for cell selection (relevant for Figure S2, S3, S5)

To estimate the chance levels for cell selection, we repeated above procedures for selection of visual category, memory selective, and choice cells after randomly scrambling the order of the labels determining the category membership being selected for (yes/no response, visual category, and new/old ground truth, respectively). We repeated this procedure 1000 times.

### 13. Single-cell decoding (relevant for Figure S5)

Single-cell decoding was done using a Poisson naïve-bayes decoder. The features used were spike counts in a 1-second window, in the interval [0.2 1.2s] relative to stimulus onset. The decoder returns the probability of a class label, given the observed spike count. The class label was binary (“yes” or “no”). The model assumes that the spike count is generated by a univariate Poisson distribution, and a separate mean rate parameter (λ) is fit to each feature-class pair. For a new observation, class membership is determined on the likelihood value. Notice that we used a single spike-count as a feature, so the naïve assumption of the decoded is no longer relevant in this case.

### 14. Population decoding (relevant for Figures 2, 3, 4, S2, S3, S4, S5, S7, S8)

Population decoding was performed on a pseudo-population assembled across sessions^68^. We present decoding results for a variety of task variables: (1) image category, (2) new vs. old, (3) choice during memory trials, (4) choice during categorization, (5) task, and (6) response type. In order to estimate the variance of the decoding performance, on each iteration of the decoder (minimum of 250 iterations), we randomly selected 75% of the cell population that was being analyzed. For example, to measure choice decoding in MFC (as shown in **Fig. 4**), we would randomly select 575/767 cells on each iteration of the decoder. The total number of available cells depended on the variable that was being decoded. For example, for response type decoding, the number of cells in MFC was 593, since 28/33 sessions included both response types. We measured decoding performance in one of two ways: (1) on a trial-by-trial basis, or (2) using condition averages. For trial-by-trial decoding, we matched the number of trials per condition contributed by each cell that was selected to participate in the population decoding. For most task variables (image category, new/old, context, effector-type) the number of samples from each cell was equal since the task structure remained the same across all subjects and sessions. For choice decoding, however, the number of instances varied, since the subjects were free to respond with a “yes” or a “no” for each stimulus. We therefore matched the numbers for the smallest group across all subjects.

In the cases where we used condition averages, the “trials” that were used to train and test the decoder were average firing rates computed on subsets of trials available for that cell. The trials that were averaged together to create a pseudo-trial, were always unique (i.e. we never resampled trials). To illustrate this procedure, consider the decoding results presented in Figure 3, which shows decoding and cross-task generalization of new/old and image category. The combination of task type, image category, new/old, and effector type results in a total of 2•4•2•2 =32 possible combinations (which contain non-overlapping sets of trials). For each cell, we computed the average firing rate for each of the 32 possible combinations. These 32 estimates then serve as the “trials” on which we train and test the decoder. For example, for decoding new vs. old, there are 16 training samples for “new” and 16 for “old”. The advantage of using condition averages is two-fold: (1) the firing rate of a cell for a given condition is more robustly estimated, (2) it eliminates biases related to imbalances of varying number of trials available for each cell and condition. The disadvantage is that trial-by-trial variability is lost. Only the following decoding results were computed on condition averages: **Fig. 3C-D, G-H, S4A-B**. As a control, we re-produce these results also with single-trial decoding (shown in Fig. S4D-E). All other decoding reported is for training and testing on individual trials.

A series of pre-processing steps were carried out before training the decoder. Firing rates for each cell were first de-trended (to account for any drift in the baseline-firing rate) and then normalized (z-scored) using the mean and standard deviation estimated from the training set. We then performed 10-fold cross validation using a linear support vector machine (SVM) decoder to estimate performance, as implemented by the ‘*fitcecoc’* function in MATLAB. We used an SVM with a linear kernel and a scale of 1. Decoding results are reported either as a function of time or in a fixed time window. Time-resolved decoding was done on spike counts measured in 500ms moving window, with a 16ms step size. For fixed-window decoding, we used spike counts in a 1-second window. The location of the window depended on the analysis we wanted to do. In **Figure 2**, for example, we used a [-1, 0] relative to stimulus onset for task type and response type decoding. In **Figure 3**, we used spike counts in [0.2, 1.2] relative to stimulus onset, for decoding image category and new/old.

### 15. Null models for testing significance of decoding performance

Throughout the manuscript, we compare the performance of our decoders against the 95^th^ percentile of a null distribution. The way that this null distribution is generated depends on the variable being decoded. For variables such as image category, new vs. old, and response (i.e. yes vs. no), we used a simple shuffling procedure for the labels. For variables such as task-type, which had structure over time (memory blocks were always preceded by categorization block), small drifts in firing rate (that was not removed by the de-trending procedure described above) might lead to inflated decoding accuracy. Therefore, for such variables, the shuffling of the variables was done in such a way as to preserve their temporal relationship. Specifically, we offset (i.e. circular shift) the labels by a random integer value (sampled from the range ±10 – 20 trials). In the case of task decoding from the baseline firing rate, this is a very conservative measure of the null decoding performance since many trials retain their original label, thereby inflating the accuracy. This also means that the mean performance of the null distribution will not be the theoretical chancel level. In the case of task decoding, the theoretical chance level is 50% (binary classification). Using the circular shift method for scrambling labels, the mean of the null distribution was ∼60%. To compare the performance between different decoders, for example MTL vs. MFC, we constructed an empirical null distribution from the pairwise differences in decoding accuracy estimated using the shuffled trial labels. To use the case of the task decoder as an example, we estimate the null distribution of the MFC decoder by shuffling the labels at least 250 times and estimate the performance for each shuffle. We do the same for the MTL decoder. We then compute the pairwise differences between the 250 values from the MFC decoder and the MTL decoder, for a total of 62500 pairwise differences. We can then estimate the likelihood of occurrence of the true difference in performance vs. this empirical null distribution. The number 250 is a representative value here and generally served as a minimum number of iterations. For lower p-values, more iterations were used.

### 16. Multidimensional scaling (relevant for Figures 3, 4)

Multidimensional scaling (MDS) was used only for visualization. We computed MDS using Euclidean distances (MATLAB function *mdscale*) on z-scored spike count data in the [0.2 1.2s] window relative to image onset. In **Figure 3E**, for example, MDS was computed on the activity across the entire population of MTL and MFC cells, averaged across the 8 conditions plotted (new/old ⨂ task ⨂ image category, where ⨂ denotes the Cartesian product). Here the image category was restricted to images of human faces and fruits, for visualization purposes. For the cross-condition generalization performance, we use all four image categories. In **Figure 4D** we compute MDS on the population of MFC cells, averaged across 8 conditions (response ⨂ task ⨂ effector, where ⨂ denotes the Cartesian product). In all cases, we use MDS to map the neural activity to a 3-dimensional space.

### 17. Normalized weight metric (relevant for Figures 4, 5, S7, S8)

The normalized weight metric is computed from the weight that a decoder assigns to a particular cell for a given classification. This weight is denoted as 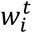, where the index ***i*** denotes the cell and the index ***t***, denotes the condition (for example, categorization or memory). The weight is converted into a normalized measure called an importance index, defined as:

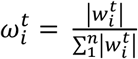

### 18. *State-space analysis (*relevant for Figure 4I)

We used Gaussian Process Factor Analysis (GPFA ^38^) to analyze the dynamics of the average population activity for the 8 conditions arising from the combination of choice (yes, no), response modality (button press, saccade), and task (memory, categorization). The recovered latent space was 8 dimensional and all similarity measurements between trajectories were performed in this space (not in the 3D projections shown in the figure). The activity was binned using 20ms windows. All analysis was computed and visualized using the DataHigh ^69^ MATLAB toolbox. Similarity measurements between two conditions were computed and averaged over the first 500ms after stimulus onset as follows:

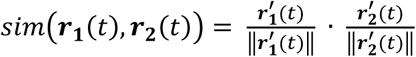

where ***r***_**1**_(*t*) and ***r***_**2**_(*t*) are the 8D state-space trajectories for condition 1 and 2 respectively.

### 19. *Spike-field coherence analysis (*relevant for Figure 5)

#### a. LFP preprocessing

The local-field potential recordings were highpass filtered at 1Hz. The raw recordings, sampled at 32kHz, were then downsampled to 500Hz. The downsampling procedure was done with the *‘resample’* command in MATLAB, which applies the appropriate antialiasing filter prior to reducing the sampling rate. For each session, we screened all MFC and MTL electrodes in order to make sure that there were no artifacts that could contaminate the spike-field metrics. We excluded all electrodes with interectal discharges (IEDs) visible in the raw trace (by visual inspection). Specifically, in screening for IEDs, we looked for large stereotyped, recurring transients in the raw recording that did not correspond to cellular spiking activity. The presence of such transients would disqualify an electrode from further consideration.

#### b. Spike-field coherence (SFC)

All spike-field coherence analysis was performed on snippets of the LFP extracted around the spike. We extract snippets for every cell-electrode pair. For example, to measure inter-area SFC between a single cell in preSMA and MTL LFPs, we extracted n snippets each (n = number of spikes) from each of the 8 ipsilateral electrodes in hippocampus and 8 electrodes in the ipsilateral amygdala. For sessions where we used a local reference (i.e. bipolar referencing), we exclude the reference wire. For intra-area coherence (ex. MTL spikes to MTL field) we also exclude the wire on which the cell was recorded to avoid contamination by spike waveform. For each snippet and for each cell-electrode pair, we compute the spike-triggered spectrum using the *FieldTrip ‘mtmconvol’* method, which computes the Fourier spectrum of the LFP around the spikes using convolution of the complete LFP traces. The spectrum was computed with a single ‘*hanning’* taper, at 56 logarithmically spaced frequencies ranging from 2 Hz on the low end, to 125 Hz on the high end. The length of the snippet window was dynamic as a function of the frequency examined; the snipped length was set to equal to two cycles of the underlying frequency at which the spectrum was estimated (i.e. 2 Hz → 2 s snippet). We estimated the phase for each snippet and for each of the 56 frequencies from the complex-valued Fourier coefficients (i.e. phasor). We use the pair-wise phase consistency (PPC) metric as the measure of coherence. For the spike-triggered power, we compute the magnitude of the spectral coefficients returned by the Fourier transform (also computed for each cell-electrode pair) for each snippet and averaged the spectra. Unless otherwise stated, all SFC results in the paper are based on spikes recorded during the baseline period between trials (1s window preceding stimulus onset).

#### c. Group comparisons using the SFC metric

When comparing two or more groups using PPC (such as memory vs. categorization), we balanced the number of spikes between the two groups. To reduce bias involved in subsampling the larger group, we resampled the spikes from the two groups 200 times, and computed the PPC metric on each iteration. The final coherence measure for a given cell-electrode pair was an average across all 200 iterations.

### 20. *Pupilometry* (relevant for Figure S1)

To test whether levels of engagement and arousal varied between tasks, we used pupillometry (pupil size; see **Figure S1J** for an example session). We compared two metrics, the baseline pupil size (0-100ms after stimulus onset, **Figure S1K**) and the slope of the pupil as it responds to the stimulus on the screen (measured from 350-600ms, **Figure S1L**). Neither metric showed a significant modulation as a function of task (p = 0.12 and p = 0.11 for size and slope respectively, sign test), thereby indicating that levels of arousal were similar for the two tasks. This analysis is based on 25 of the 28 sessions where we measured eye movements. The remaining 3 sessions were not used because the measurement of the pupil was determined to be too noisy.

## Data availability

The data that support the findings of this study are available on reasonable request from the corresponding author.

## Code availability

We used the following toolboxes, all of which are available as open source: OSort for pre-processing and spike sorting, FieldTrip, DataHigh, CircStat, and Psychtoolbox. All other work was done using standard built-in MATLAB functionality.

## Supplementary Figures

**Supplementary Figure 1.**
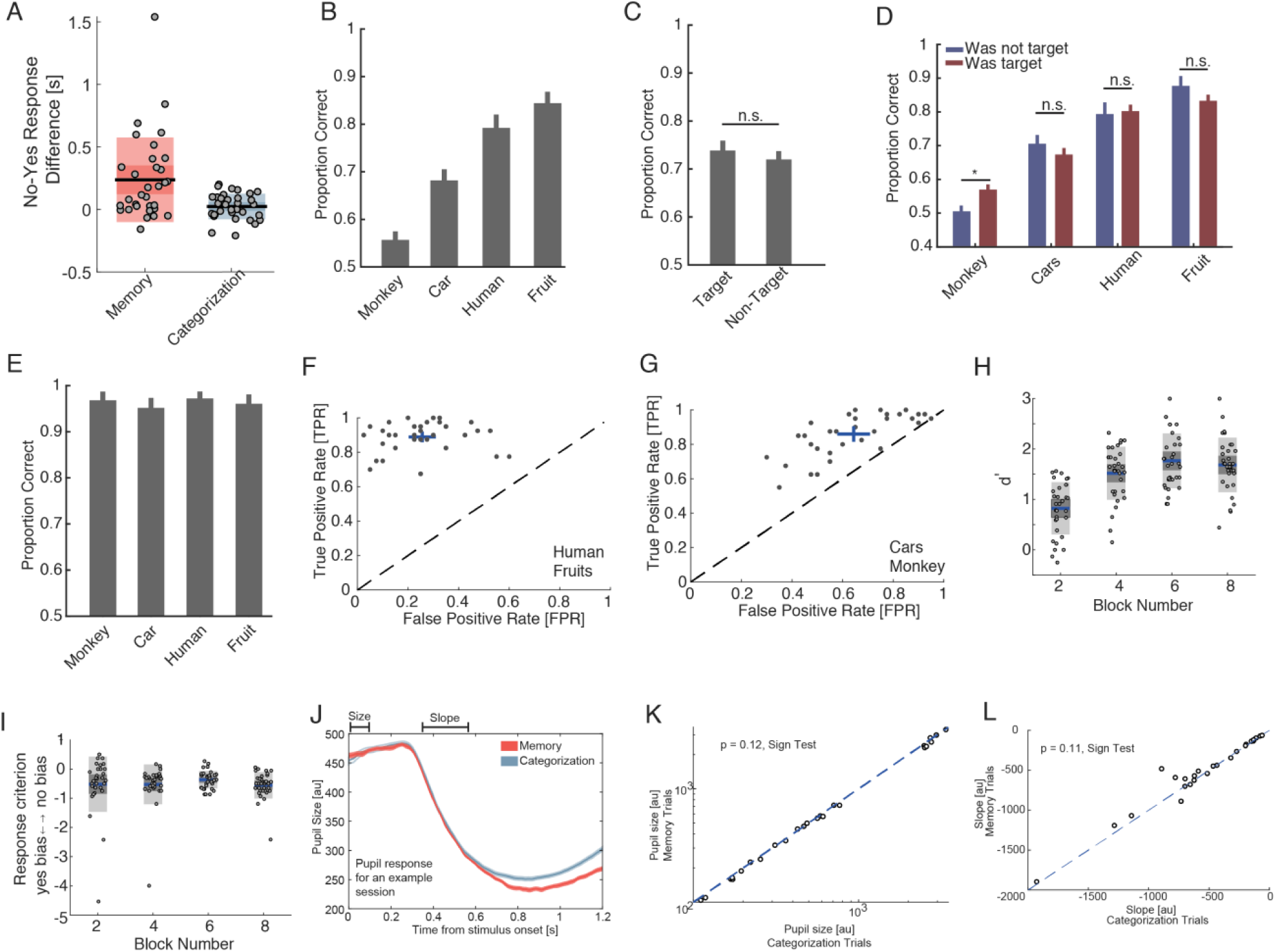
Additional behavioral results. **(a)** RT difference between “yes” and “no” responses for both tasks. RTs were significantly different for the memory task (p=3.4e-4; as expected from a declarative memory task ^9^ but not for the categorization task (p=0.21) task (t-test, light shading indicates ± std, whereas darker shading indicates ± sem). **(b)** Performance on memory trials varied as a function of image category (1×4 ANOVA, p = 3.45e-19). **(c)** Making an image category the target on a categorization block did not significantly change recognition accuracy for that category on follow-up memory blocks (p = 0.97, t-test). **(d)** Same as (c) but shown separately for each category. Recognition performance increased significantly after being a target only for the monkey category (p = 0.02, t-test; uncorrected). (**e**) Performance on the categorization trials did not significantly dependent on image category (1×4 ANOVA, p = 0.74). (**f**) ROC analysis of the performance on memory trials for the two best image categories (human faces and fruits). (**g**) ROC analysis of the performance on memory trials for the two difficult image categories (cars and monkeys). While the true positive rate (TPR) is not different between these two image groups, (p = 0.24, t-test), the subjects produced significantly more false positives for the more difficult group (p=1.2e-13, t-test). **(h)** d’ as a function of block number (4 memory blocks). d’ increased significantly across blocks (1×4 ANOVA, p = 2.6e-11). **(i)** The response bias for each subject as a function of block number. The response bias did not vary significantly across blocks (1×4 ANOVA, p = 0.6). **(j)** Example pupil response during a session. We focus on two measures: (1) the baseline size, measured shortly after image onset, and (2) the slow of the pupil response. The differences that emerge after 600ms are due to response time differences between the tasks (the image goes off the screen earlier for categorization trials). **(k)** Scatter of the pupil size, measured shortly after image onset, for all sessions with stable pupilometry (25/28 sessions with eye tracking). There was no significant difference between tasks (p = 0.12, sign test). **(l)** Scatter of the slope size across sessions with good pupilometry (25/28 sessions with eye tracking). There was no significant difference between the tasks (p = 0.11, sign test).

**Supplementary Figure 2.**
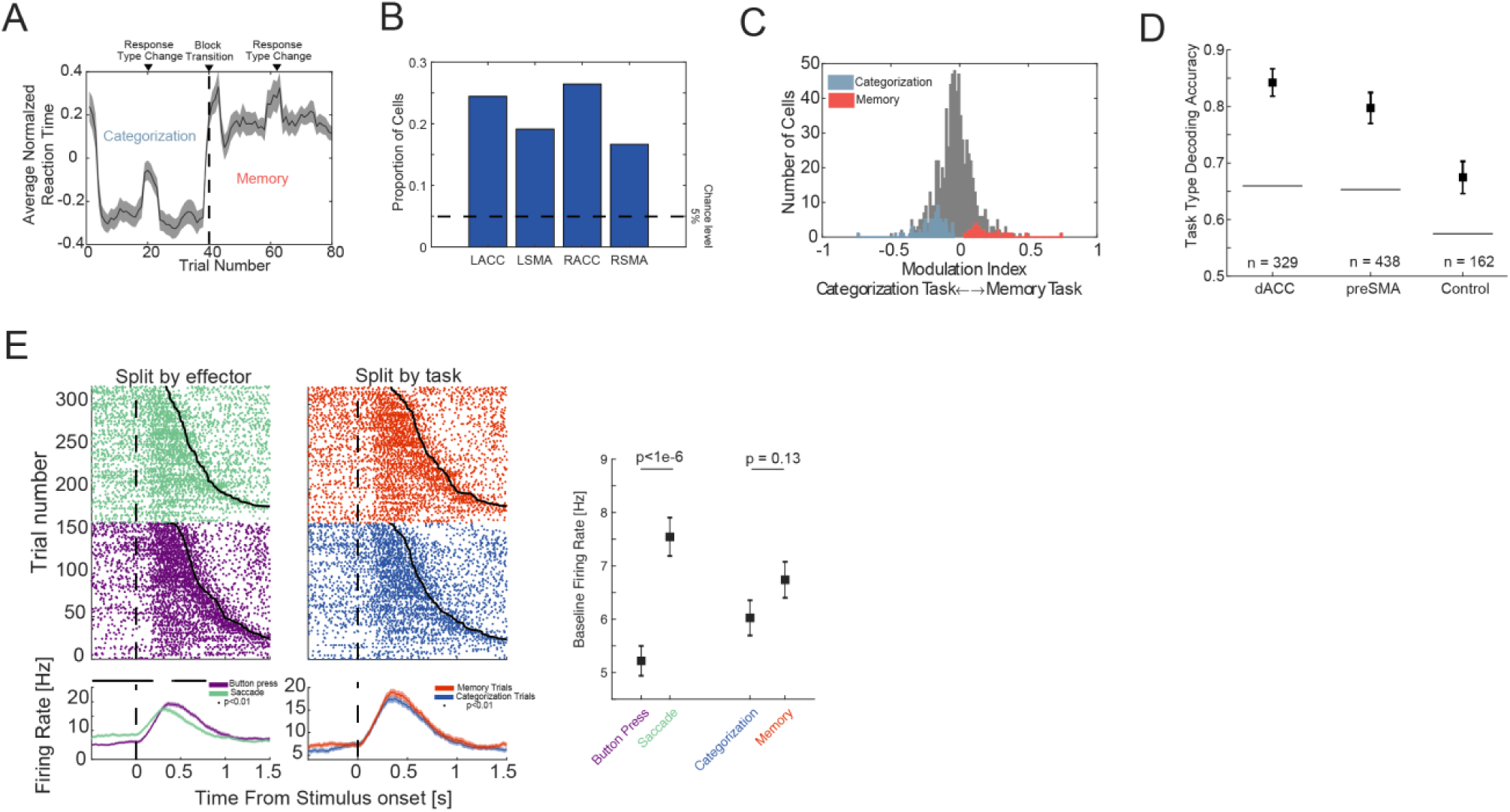
Context information in the medial frontal cortex. **(a)** Average reaction time as a function of trial number, averaged across all subjects and block switches. Trial 41 marks the transition from a categorization task to a memory task. Halfway through each block, there is a change in response modality. Reaction time is smoothed with a 5-trial kernel. **(b)** The proportion of cells sensitive to either task type or response modality, as a function of hemisphere and area in the medial frontal cortex (L=left, R=right). The proportion of cells sensitive to context during the baseline period was significantly greater in the dACC than pre-SMA (χ^2^ test of proportions, p = 0.02). **c)** Selected cells are sorted into two groups, memory task-preferring or categorization task –preferring based on their firing rate during the baseline period. Shown here is a histogram of the modulation index, 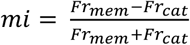. **(d)** Context can be decoded from both the dACC and pre-SMA. For comparison shown is decoding accuracy for the control sessions (in which the response time does not differ between task types). The numbers indicate the number of cells that were included for each of the three decoders. Bars are standard deviation of decoding accuracy across all iterations of the fitting procedure (n = 250). The dotted lines mark the 95^th^ percentile of the null distribution of decoding accuracy, computed by shuffling the labels, and performing the fit 250 times. (**e**) An example cell recorded in the pre-SMA that shows baseline modulation of firing rate with effector type (left side) but not task type (right side).

**Supplementary Figure 3.**
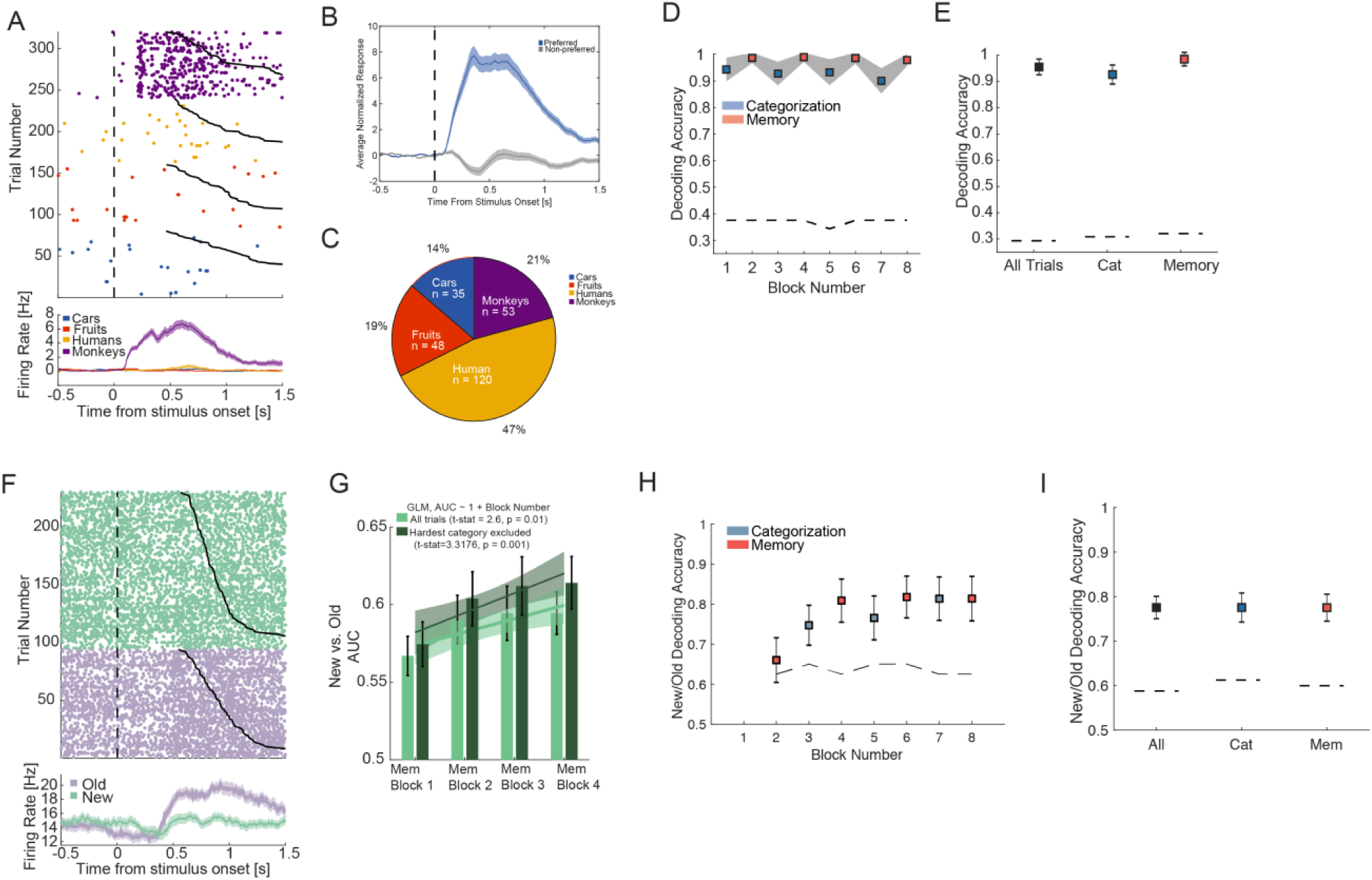
Visually-and memory selective MTL cells are not sensitive to task demands. **(a)** Example visually selective cell recorded in the amygdala. The cell responds selectively to images of monkey faces with little activity for the other three image categories. **(b)** Average normalized response to preferred (blue) vs. non-preferred images for all visually selective cells in the MTL (n=264/663, see methods for selection criteria). **(c)** Breakdown of the preferred category of visually selective cells in the MTL. As previously reported, most cells respond to faces of conspecifics. **(d)** Trial-by-trial decoding of image category over the 8 blocks within a session. The gray shading indicates the standard deviation across 200 iterations of the population decoder (see methods), using the [0.2 1.2s] time bin after stimulus onset. The decoder was trained using all trials. Shown here is the cross-validated accuracy of the decoder on each block separately, with categorization blocks in blue and memory blocks in red. The dotted black line shows the 95^th^ percentile of the null distribution, computed by shuffling the labels. The chance level is 25%. **(e)** Same as in (d) but collapsed across task types. The dotted lines once again indicated the standard deviation across 200 iterations of the decoder, using different subsets of cells and trials (see methods of details). **(f)** Example memory selective cell in the MTL. **(g)** Average AUC across all memory selective cells (n=73/663, see methods for model used to identify this cell type) for new vs. old stimuli, shown across all memory blocks. The number of new and old stimuli in each block is equal (20 of each). In light green, we show average AUC across all cells, for all the 4 image categories. New and old stimuli became more separable over the blocks (GLM, AUC ∼ 1 + Block_Number, t-stat = 2.6, p = 0.01). If the hardest category is removed (monkey images), the effect becomes more evident (t-stat = 3.32, p = 0.001). **(h)** Trial-by-trial population decoding of new vs. old across all blocks in the session. The first block is excluded because it does not contain “old” stimuli. The dotted line shows the 95^th^ percentile of the null distribution of decoding performance (chance level is 50%). **(i)** Same as (f) but collapsed across task type.

**Supplementary Figure 4.**
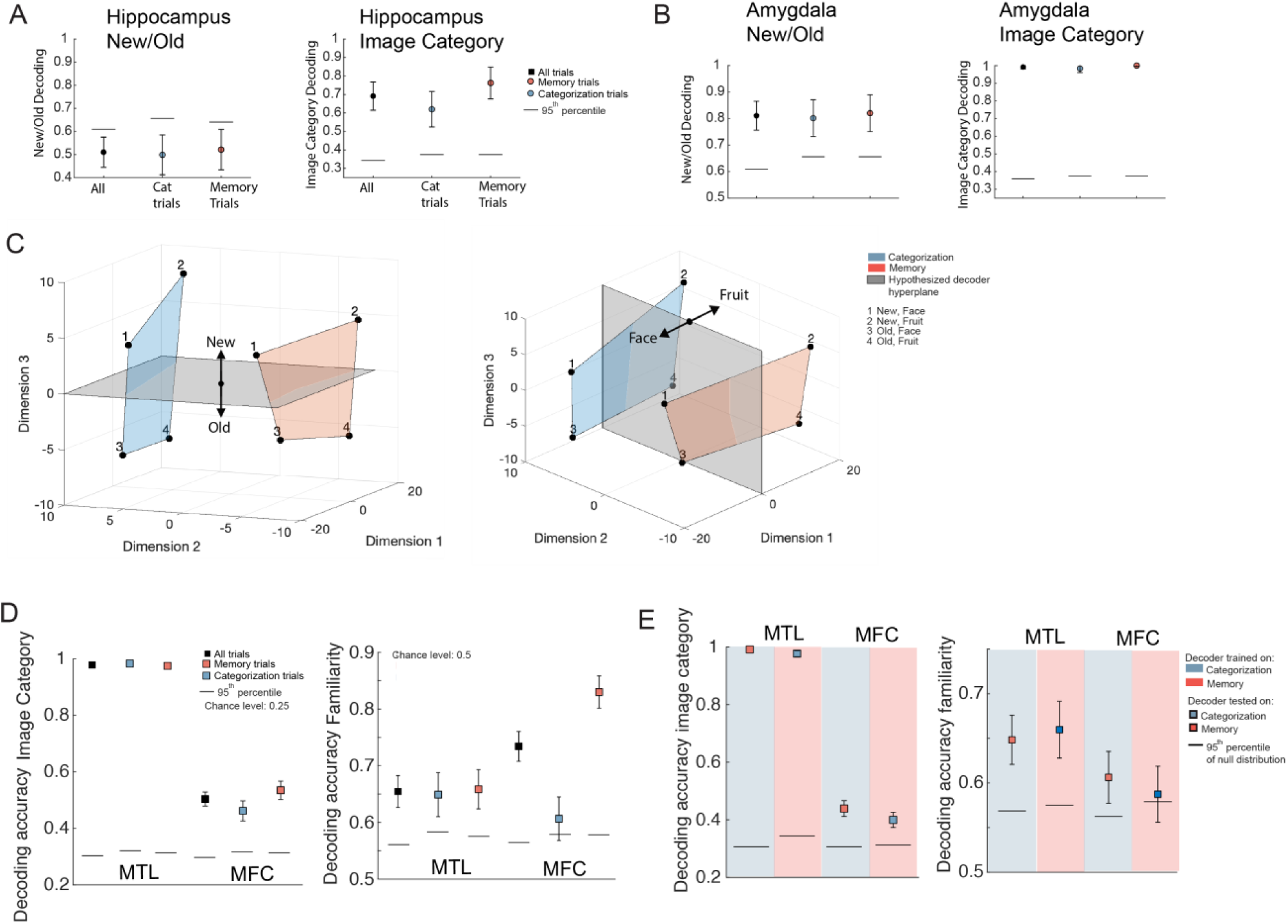
Cross-task generalization of new/old and image decoding across the four brain areas studied. **(a-b)** Decoding accuracy in (a) hippocampus, (b) amygdala of new/old (left column) and image category (right column). Each decoder was trained on condition averages (see methods). We show the performance separately for all trials, categorization trials, and memory trials. (**c**) Rotated version of the MDS plots shown in **Fig. 3E**, with *example* decision boundaries for a new/old (left) and image category (right) decoder. The locations of the condition averages are computed from the population activity in the MTL, whereas the decision boundary is schematized to show an example decoder that would generalize well across tasks. (**d**) We reproduce the results shown in **Fig. 3C-D** with single-trial training/testing. (**e**) Cross-task generalization of image category (left) and familiarity (right) as shown in **Fig. 3G-H**, reproduced using single-trial training/testing.

**Supplementary Figure 5.**
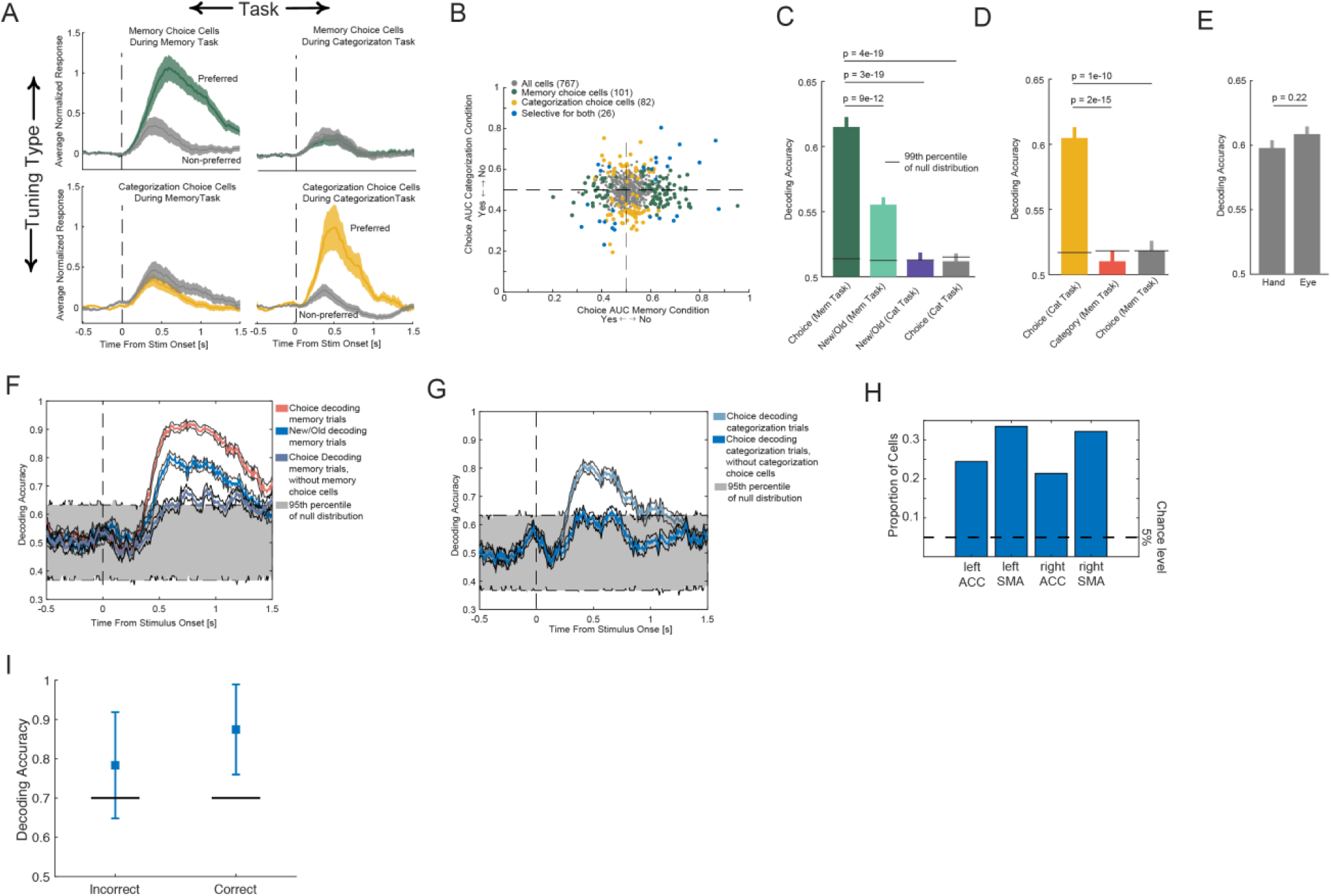
Choice cells in medial frontal cortex are sensitive to task demands – additional single-cell analysis. (**a**) Average PSTHs for memory choice cells (green, n = 101/767) and categorization choice cells (yellow, n = 82/767), shown separately for the two tasks. Memory and categorization choice cells were selected independently using trials from the corresponding task (see methods for selection model). Omitted from this visualization are choice cells that were selected in both task conditions (n=26/767). (**b**) Population summary. AUC values were computed separately for response made during the memory and categorization condition. A negative AUC value indicates a preference for “yes” responses and a positive one indicates a preference for “no” responses. Yellow indicates categorization choice cells, green indicates memory choice cells, and purple indicates cells that signal choice in either task. **(c)** Single cell decoding across all memory choice cells (101/767). Decoding performance is shown for choice during the memory trials (green), new vs. old during the memory trials (cyan), new vs. old during the categorization trials (purple), and choice during the categorization trials (yellow). **(d)** Single cell decoding across all categorization choice cells (82/767). Decoding performance is shown for choice during the categorization trials (yellow), image category during the memory trials (orange), and choice during the memory trials (gray). (**e**) Comparison of choice decoding (collapsed across both tasks) performance between response modalities. There was no significant difference. (**f-g**) Population decoding performance as a function of time during the memory (g) and categorization (h) task. Performance was reduced significantly after choice cells were removed from the population. **(h)** Proportion of selected choice cells in medial frontal cortex, separated by area and hemisphere. The proportion of choice cells found is greater in the pre-SMA than dACC (χ^2^ comparison of proportions, p = 0.004). (**i**) Trial-by-trial choice decoding at the population level. The decoder was trained on equal examples from the following memory trials: (1) yes-correct, (2) yes-incorrect, (3) no-correct, (4) no-incorrect. Cells were included in the analysis only if there were at least 10 instances of each of the four trial types (n=347 cells). Choice can be decoded from the MFC population both for correct and incorrect trials.

**Supplementary Figure 6:**
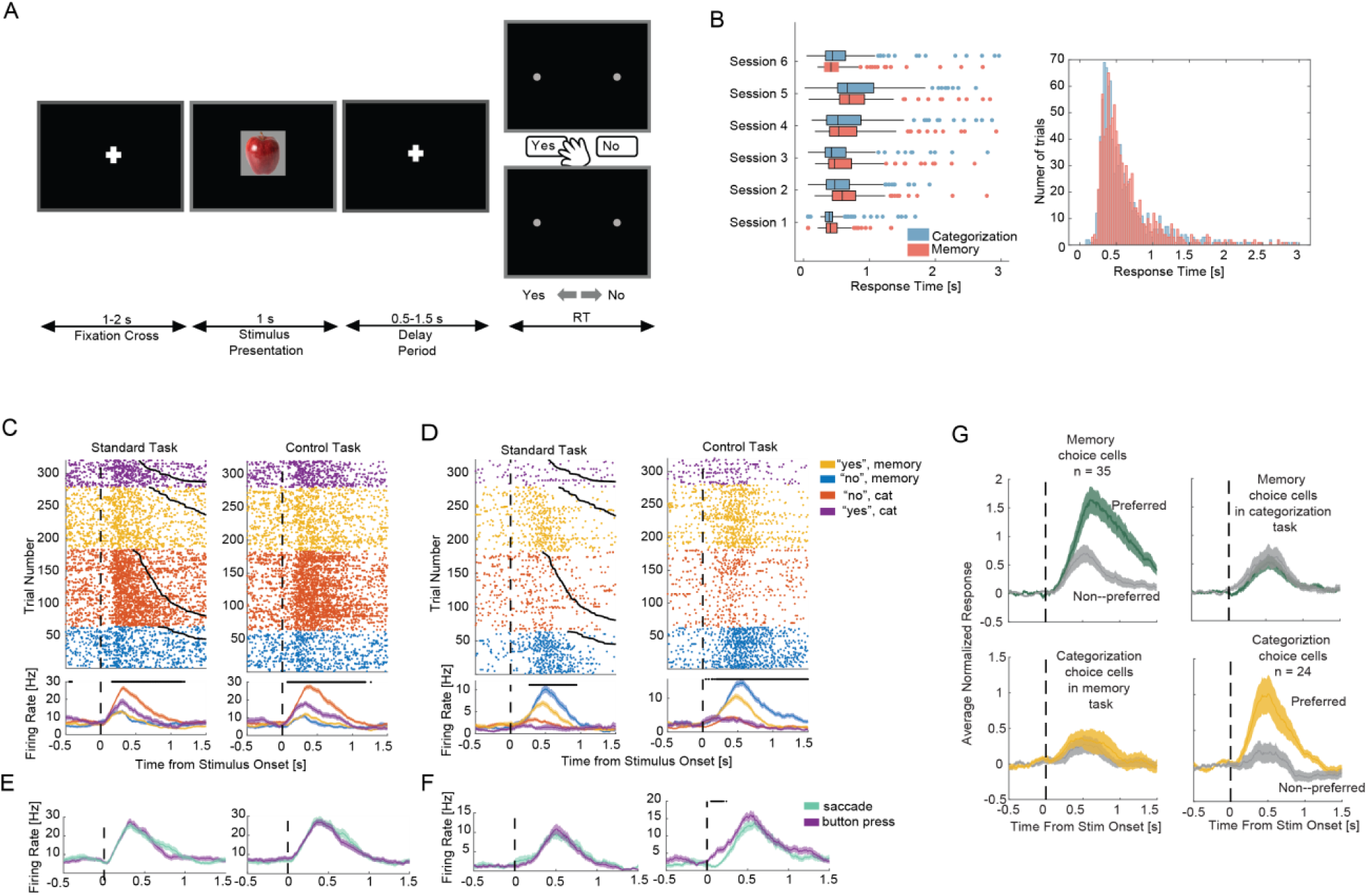
Choice signals during a non-reaction time control task. **(a)** Task layout for the non-reaction time control task. Subjects are instructed to wait until the response screen comes up before registering their response with a button-press or a saccade. The stimulus length is fixed at 1 second, for both the categorization and memory trials. **(b)** The response times between the categorization and memory trials are no longer different (mean ± std, 0.67 ± 0.57s and 0.72 ± 0.77s for categorization and memory trials respectively, p = 0.1, 2-sample t-test). **(c-d)** Raster plots and PSTH of two example choice cells recorded in the dACC (C) and pre-SMA (D) during the standard task (left panel) and control task (right panel). Notice that there is no button press or saccade prior to 1.5 second during the control task. **(e-f)** The preferred response for the cell shown in C (“no” during categorization) and the cell shown in D (“no” during memory condition) split up by effector type, with saccade responses in green and button press in purple. **(g)** Average PSTH for preferred and non-preferred responses across all the choice cells identified in the control task. Top row shows the preferred vs. non-preferred response of memory choice cells during the memory task (left panel) and categorization task (right panel). The same is shown for categorization choice cells in the bottom row.

**Supplementary Figure 7.**
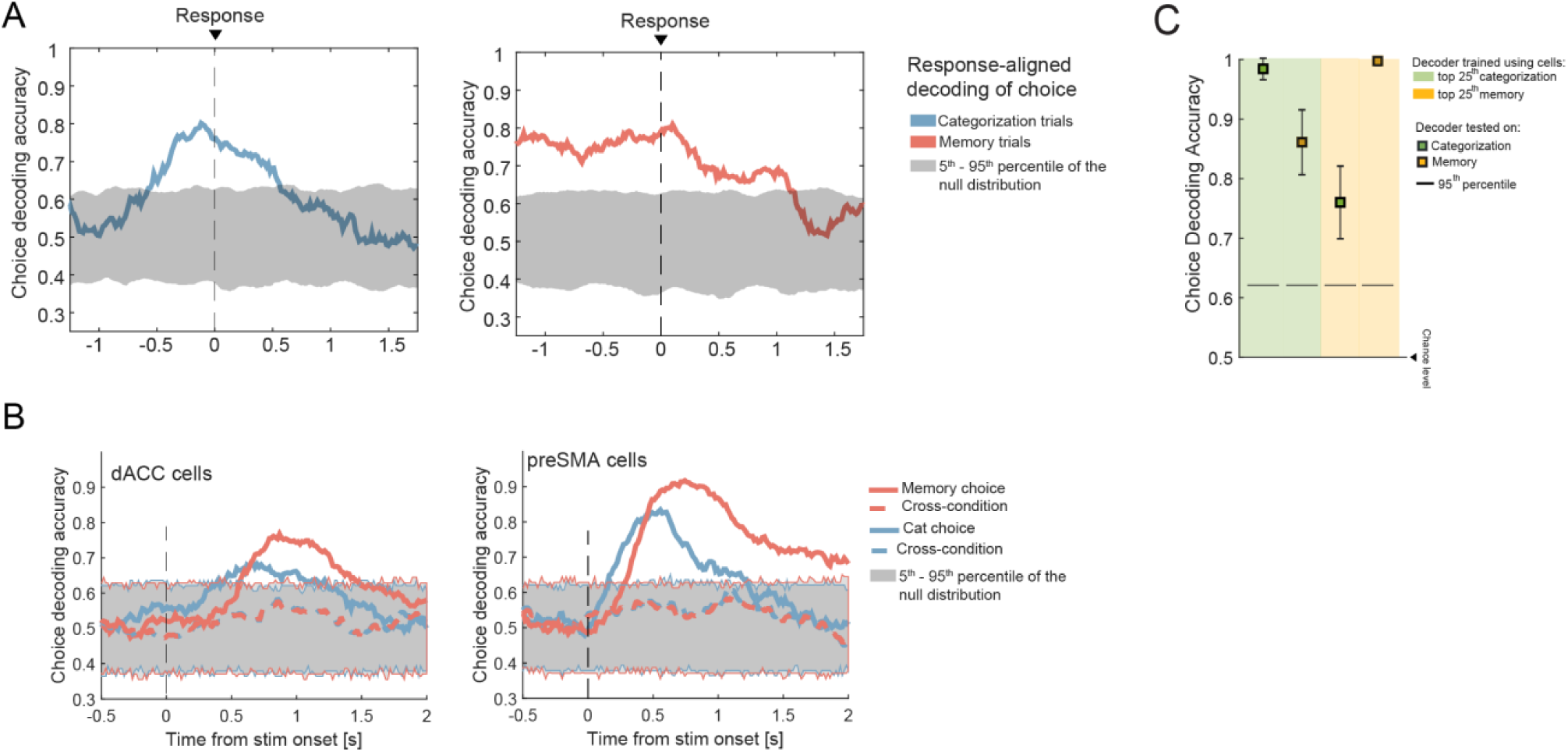
Cross-task generalization of choice signals by area. Population-level decoding of choice during the categorization trials (left) and memory trials (right) using firing rates that are aligned to the response time instead of stimulus onset. Compare with **Figure 4e. (b)** Cross-task generalization of choice decoding in the dACC (left) and pre-SMA (right) shown as a function of time. This is the same analysis as that in Figure 4e, but shown separately for the two areas. **(c)** The cells that are in the top 25^th^ percentile of the weight index distribution for either task (see **Fig. 4I**) can be used to train a *new* decoder that accurately predicts choice in the other task, albeit with a significantly diminished performance. Note that this is ***not*** cross-condition generalizations since we are training a new decoder on a subset of the MFC cells.

**Supplementary Figure 8:**
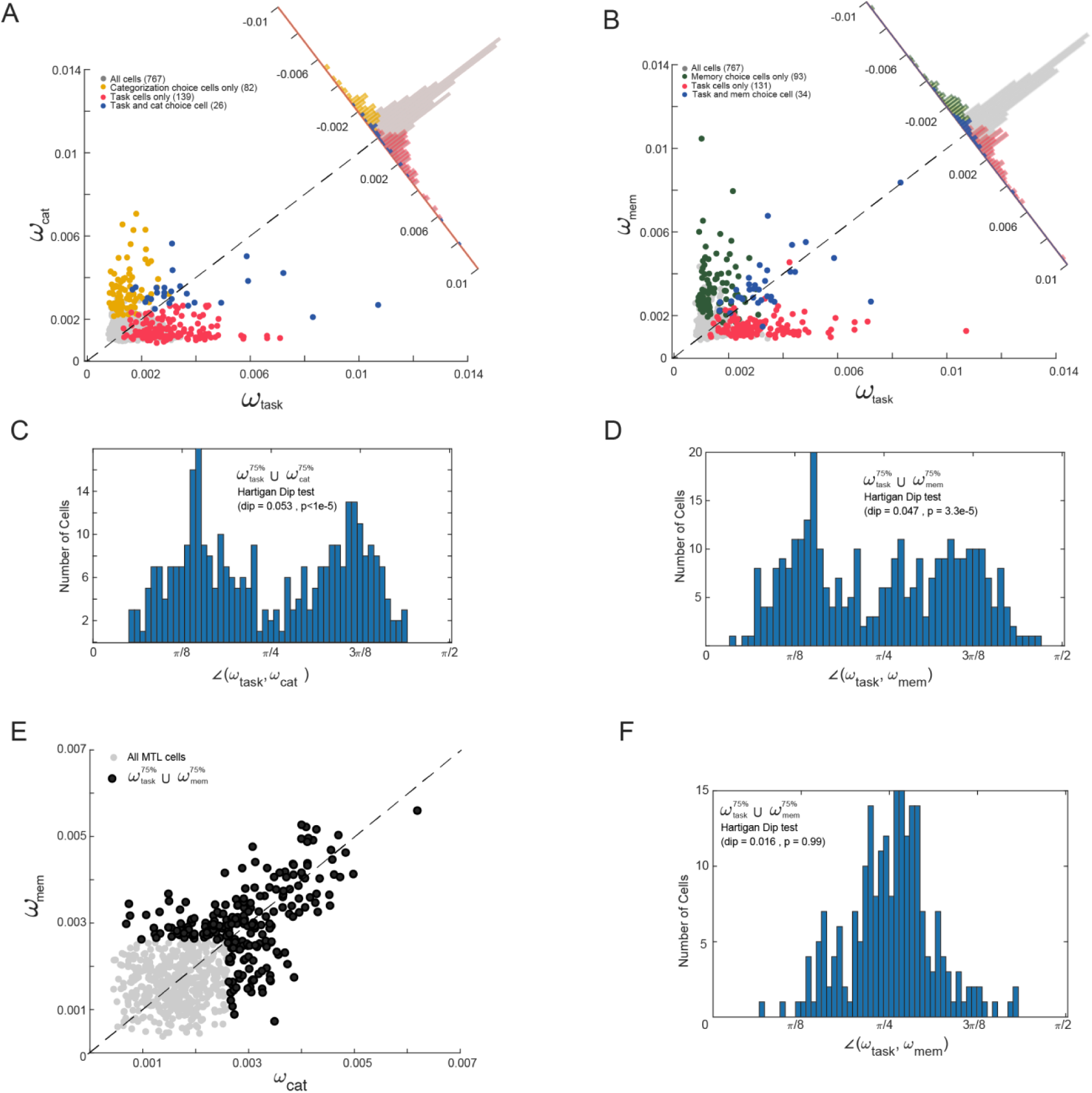
Comparison of task and choice cells using assigned decoder weight. **(a)** Scatter plot of the weight assigned by a decoder to each cell in decoding categorization choice (y-axis), and task (x-axis). The features for the choice decoder are firing rates across the entire MFC population in the [0.2 1.2s] window after stimulus onset. The features for the task decoder are firing rates computed during the pre-stimulus baseline period, [-1 0s] with respect to image onset. As in Figure S6, the decoder weight is converted into a normalized measured (importance index). Superimposed are the populations of categorization choice cells and task cells, as identified by the choice and task selection models described in the Methods section. **(b)** Same as in (a), but shown for memory choice decoding and task decoding. Highlighted in green are the memory choice cells, and in pink are the task cells (see Methods for selection model). The cells that qualify as both are shown in blue. (**c**) Similar to Figure S6D, we look at the cells that have a high weight index for either categorization-choice or task decoding. Specifically, we take the *union* of the sets of cells whose weight index is in the top 25^th^ percentile for either task or categorization-choice decoding. For these cells, we plot the angle created by the vector 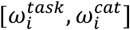 with respect to the x-axis (i.e. the task axis). We test for bimodality with a Hartigan dip test (dip = 0.053, p<1e-5), the result of which suggests that these are largely different populations of cells. (**d**) Same as (c) but in this case we measure the overlap between memory choice cells and task cells. The histogram shows two modes, suggesting non-overlapping populations of cells (dip = 0.047, p = 3.3e-5). (**e**) Decoding of image category from the MTL population is a good example of a case where the same cells are recruited for decoding in the memory and categorization task. Shown in light gray is the weight index for all MTL cells, computed separately for the categorization and memory task. The dark dots indicate the union of the sets of cells that have a weight index top 25^th^ percentile for either task. (**f**) Hartigan dip test for the weight index pair assigned to each cell in black from (e). The distribution is centered at π/4, which suggests that the same cells are recruited to decode image category during the memory and categorization tasks (dip = 0.016, p = 0.99).

**Supplementary Figure 9:**
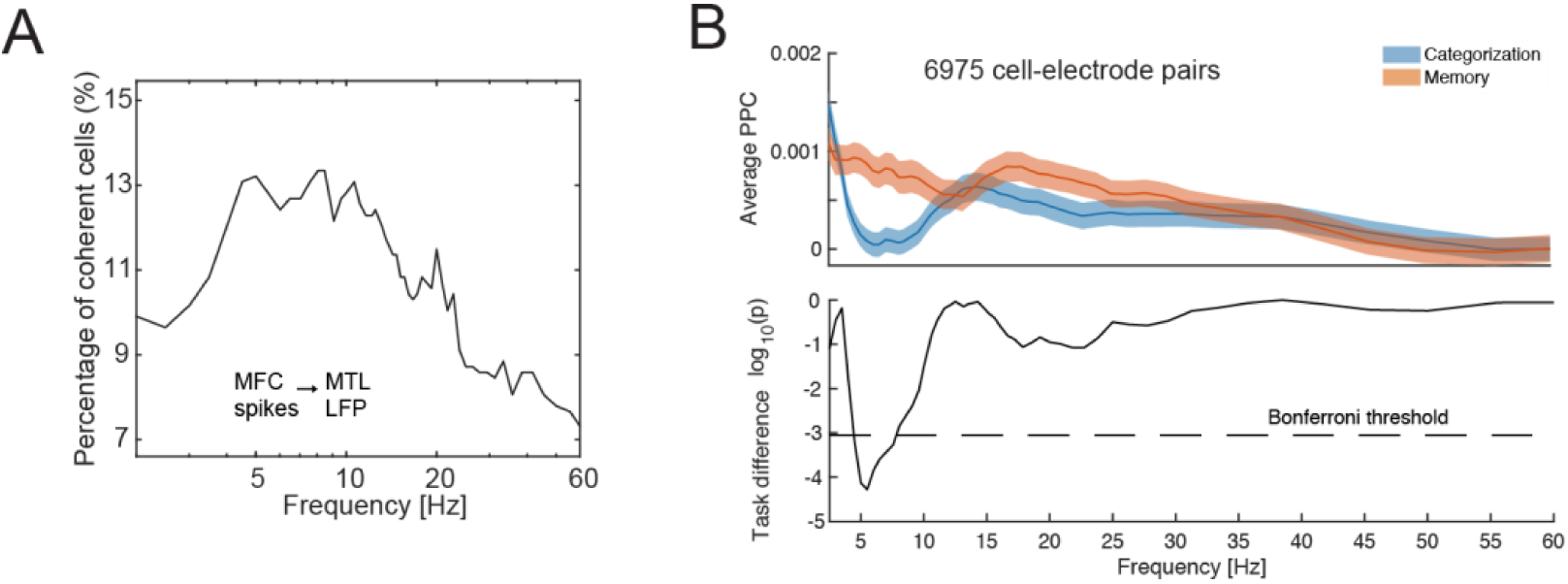
Controls for inter-area spike filed coherence between MFC cells and MTL local filed potential. **(a)** Proportion of MFC cells that are coherent with hippocampal oscillations using spikes from the inter-trial period of all trials. Coherence was determined using the Rayleigh test for non-uniformity of a circular distribution. Since the comparison was done across many electrodes (N_channels_ can be anywhere from 0 to 16, depending on the number of LFP recordings accepted after screening for artifacts, see **Methods**), the significance threshold was corrected appropriately for multiple comparisons using FDR (false discovery rate). **(b)** Same as **Fig. 5C**, with task cells removed (n=165/767). A cell was labeled as a task cell (see **Fig. S2** and **Methods** for selection) if it should significant modulation of firing rate as a function of task type.

**Table S1.** Recording sessions.

| Patient ID | Session # | # MTL cells | # MFC cells | Response modality used (1=button press only, 2 eye+hand) |
| --- | --- | --- | --- | --- |
| P41 | 1 | 1 | 5 | 2 |
| P41 | 2 | 3 | 9 | 2 |
| P41 | 3 | 2 | 1 | 2 |
| P42 | 4 | 13 | 32 | 1 |
| P42 | 5 | 20 | 42 | 1 |
| P43 | 6 | 19 | 0 | 2 |
| P43 | 7 | 25 | 1 | 1 |
| P43 | 8 | 23 | 0 | 2 |
| P44 | 9 | 11 | 59 | 1 |
| P44 | 10 | 8 | 40 | 1 |
| P47 | 11 | 28 | 8 | 2 |
| P47 | 12 | 37 | 8 | 2 |
| P47 | 13 | 33 | 5 | 2 |
| P48 | 14 | 19 | 39 | 2 |
| P49 | 15 | 2 | 3 | 2 |
| P49 | 16 | 5 | 2 | 2 |
| P51 | 17 | 20 | 38 | 2 |
| P51 | 18 | 20 | 18 | 2 |
| P51 | 19 | 18 | 21 | 2 |
| P51 | 20 | 18 | 21 | 2 |
| P51 | 21 | 11 | 14 | 2 |
| P53 | 22 | 8 | 12 | 2 |
| P53 | 23 | 16 | 21 | 2 |
| P56 | 24 | 32 | 14 | 2 |
| P56 | 25 | 15 | 11 | 2 |
| P56 | 26 | 34 | 6 | 2 |
| P57 | 27 | 31 | 23 | 2 |
| P57 | 28 | 28 | 34 | 2 |
| P57 | 29 | 28 | 34 | 2 |
| P58 | 30 | 43 | 75 | 2 |
| P58 | 31 | 43 | 75 | 2 |
| P58 | 32 | 34 | 53 | 2 |
| P61 | 33 | 15 | 43 | 2 |
| **Total** |  | **663** | **767** | **28 with, 5 without eye tracking** |

